# Systemic ligand-mimicking bioparticles cross the blood-brain barrier and reduce growth of intracranial triple-negative breast cancer using the human epidermal growth factor receptor 3 (HER3) to mediate both routes

**DOI:** 10.1101/2021.06.07.446634

**Authors:** Felix Alonso-Valenteen, Sam Sances, HongQiang Wang, Simoun Mikhael, Jessica Sims, Michael Taguiam, Dustin Srinivas, Erik Serrano, Briana Ondatje, James Teh, Michelle Wong, Kimngan Nguyenle, Tianxin Miao, Rebecca Benhaghnazar, John Yu, Clive Svendsen, Ravinder Abrol, LK Medina Kauwe

## Abstract

Crossing the blood-brain barrier (BBB) and reaching intracranial tumors is a significant clinical challenge for targeted therapeutics and contributes to the poor prognosis for most patients with brain malignancies. Triple-negative breast cancer (TNBC) has a high propensity for metastasis to the brain and lacks cell surface markers that can be recognized by current targeted therapies used in the clinic, thus limiting therapeutic options. The human epidermal growth factor receptor HER3 (or ErbB3) has emerged as a biomarker of therapeutic resistance and metastasis in a growing range of tumor types and may serve as a possible therapeutic target for TNBC. Accordingly, we have developed HER3-targeted biological particles (bioparticles) that assume polyhedral capsid shapes when encapsulating nucleic acid cargo, forming nano-nucleocapsids (NNCs). The NNCs exhibit systemic homing to resistant and metastatic breast tumors, including TNBC, due to the high cell surface densities of HER3 on these tumors. Here we describe our discovery that HER3 is also prominently expressed on the brain endothelium and can mediate the passage of HER3-targeted NNCs across the BBB and into triple-negative breast tumors localized in the brain. Our findings show that HER3 is present at high levels on the vasculature (but not extravascular parenchyma) of both mouse and human adult brain specimens and associates with the extravasation of systemic HER3-targeted NNCs in mice and in a human model of the BBB (BBB chip). Furthermore, systemically delivered NNCs carrying tumoricidal agents reduced the growth of intracranial TNBC tumors in mice (representing metastatic breast tumors that have established in the brain) and exhibited improved therapeutic profile compared to current therapeutic interventions (liposomal doxorubicin) used in the clinic. This study addresses the major clinical problem of systemically delivering targeted therapeutics across the blood-brain barrier (BBB), and demonstrates a new route for not only accomplishing this but also for reaching tumors localized in the brain.

## INTRODUCTION

The blood-brain barrier (BBB) prevents most systemic therapeutics from entering the brain parenchyma and, more importantly, intracranial tumors, thus contributing to the poor prognoses of patients with brain malignancies. Brain metastatic breast tumors entail only a 1-month median survival of untreated patients, while whole-brain radiotherapy increases median survival ~4-6 months.^1^ Options available to these patients include surgical procedures (which pose high risk in the brain and cannot safely remove all tumor tissue), chemotherapy, and radiation therapy.^2^ Disseminated tumors and micro-metastases pose challenges for surgical removal, and untargeted drugs that can reach such tumors affect healthy tissue as well. Targeted antibody therapies that can combat peripheral tumors are unable to cross the BBB, making them ineffective against brain-localized tumors-- including secondary tumors that have metastasized to the brain-- despite the expression of respective therapeutic targets in such tumors.^3–4^ For example, the monoclonal antibody trastuzumab (Herceptin^®^), which is directed against the ErbB growth factor receptor kinase HER2, effectively targets peripheral HER2+ breast tumors but cannot transit the brain endothelium to reach HER2+ intracerebral lesions, including brain metastatic HER2+ breast cancer. ^5–7^ Triple-negative breast cancer (TNBC), which is HER2 negative and highly brain-metastatic, has even fewer options due to the lack of specific cell surface biomarkers that can be recognized by targeted therapies currently used in the clinic.^8^ Temporary opening of the BBB using osmotic regulators may transiently allow molecules into the brain non-specifically, between endothelial cells, but poses additional risk due to the non-selectivity and unregulated nature of this procedure.^9–10^ Known receptors on the brain vasculature such as GLUT-1 or transferrin receptors have been explored as routes of transfer across the endothelium;^11–13^ however, the ubiquitous expression of these receptors may reduce brain and tumor-specific targeting, and may not necessarily associate with brain-metastases. Moreover, small molecules such as the ErbB tyrosine kinase inhibitor lapatinib can diffuse across the cell membrane, yet show poor delivery and poor efficacy to brain-metastatic tumors^14–15^. It is therefore a major clinical challenge to find ways to shuttle targeted therapies across the BBB to combat malignancies in the brain.

Numerous studies in recent years have highlighted an expanding role of HER3 (or ErbB3) in tumor progression,^16–28^ showing that its increased cell surface density associates with therapeutic resistance and metastasis, including metastasis to the brain.^16–17^, ^22, 29–33^ Increased levels of cell surface HER3 have been identified on both HER2+ and triple-negative breast tumors that resist current targeted therapies used in the clinic, and metastasize to the brain.^34–35^ HER3 is increased on a broad range of other tumor types as well, including prostate, gastric, colon, lung, pancreatic, head and neck, ovarian, cervical, glioblastoma and skin cancers^18–28^ in experimental models, and patient specimens^36^. HER3 belongs to the same ErbB growth factor receptor kinase family as HER2, but it lacks innate kinase activity ^37^. Thus, despite its increased expression on malignant cells, it is a poor target for receptor kinase inhibition.

Although HER3 targeted therapies are currently lacking in the clinic, a HER3-targeted antibody-drug conjugate (ADC; U3-1402) is currently undergoing Phase II testing for metastatic breast cancer.^38^ A HER3 antibody, patritumab/U3–1287, developed for targeting non-small cell lung cancer progressed to Phase III testing but this trial has since been terminated due to predefined criteria not reached.^39–40^ Other HER3 antibodies are at various earlier stages of clinical trials for peripheral tumors and are largely aimed at ligand blocking and preventing downstream activation.^39–40^ Given the resistance of HER3+ tumors to signal-blocking therapies, including combination therapy,^39^ it is likely that alternative approaches circumventing the modulation of downstream activation would still be needed. Additionally, the restrictions imposed by the BBB on effective brain entry of monoclonal antibodies as described earlier are likely to limit an ADC approach to reaching intracerebral tumors.

We have bioengineered a tumor-invading protein, HPK (previously known as HerPBK10^41^), that can home in on breast tumors resisting trastuzumab (Tz) and other ErbB receptor family inhibitors through its high avidity targeting of HER3.^42^ Unlike antibodies, HPK uses the receptor engagement of the natural HER3 ligand, heregulin (or neuregulin)^43^ to exploit a ligand-induced route of cell entry. HPK also takes advantage of the capsid-forming and membrane penetration functions of the adenovirus-derived penton base protein to encapsulate and deposit macromolecular cargo into HER3-expressing cells.^44^ Cargo encapsulation forms biologically-based particles or bioparticles that use neuregulin mimicry to breach HER3+ tumor cells while reducing HER3 activation by wild-type neuregulins^44^. Through ligand mimicry, HPK bioparticles evade immuno-recognition by anti-adenovirus antiserum and exhibit no detectable stimulation of neutralizing antibodies in contrast to whole adenovirus in immune-competent mice.^44^

In the present study we demonstrate that HPK can form nucleic acid-containing bioparticles that not only invade HER3+ tumors but also distribute to the brain in contrast to cargo lacking neuregulin-directed delivery. We subsequently found that HER3 is prominently expressed on both mouse and human brain endothelium and associates with systemic bioparticle transfer into the brain parenchyma. The association of HER3 with metastasis to the brain ^34–35, 45^ led us to then ask whether brain-localized HER3+ tumors can be accessed by systemic HPK bioparticles. Therefore, here we test the hypothesis that HER3 is present on the BBB and TNBC, and can mediate the transport of systemic HER3-homing bioparticles across the BBB and into HER3-expressing TNBC tumors in the brain; and we evaluate the capacity for systemically delivered bioparticles carrying tumoricidal agents to reduce intracranial tumor growth compared to untargeted chemotherapy, which is among the few clinical options available for treating brain-metastatic TNBC. Here we used mice bearing intracranial implants of TNBC, which serves as a preclinical model of secondary TNBC tumors that have established in the brain.

The preponderance of studies on HER3 have focused on its contribution to tumor progression and resistance while little is known about a potential role in passage across the BBB. Neuregulins normally facilitate neuronal maturation, myelination and repair in the central and peripheral nervous system ^46^. If a route exists that mediates neuregulin passage across the BBB, such a route could be engaged by HPK bioparticles. Therefore, these studies also probe HER3 localization in disease-free mouse and human brain specimens and interrogate the contribution of HER3 to bioparticle extravasation in mice and in a human induced pluripotent stem cell (iPSC)-derived organ-on-a-chip model (BBB chip). The use of currently available mouse models of intracranial TNBC and human-derived models of the BBB enable these studies to rigorously evaluate the translational impact of HER3 targeted biocarriers on brain-localized tumors and uncover a new route for systemic delivery into the brain.

## METHODS

### Materials

PBS (50 mM NaH_2_PO_4_, 150 mM NaCl, pH 7.5); PBS++ (1% Ca^2+^ and 1% Mg^2+^ in 1X PBS); Buffer A (DMEM containing 20 mM HEPES, pH 7.4; 2 mM MgCl_2_; and 3% BSA); Human brain tissue specimens were obtained through Tissue for Research Ltd.

### Cells

All cell lines were originally obtained from the American Type Culture Collection and maintained at 37°C with 5% CO_2_ under mycoplasma-free conditions in complete medium [Dulbecco’s modified Eagle’s medium (DMEM) supplemented with 10% fetal bovine serum, 100 U/mL penicillin, and 100 μg/mL streptomycin].

### Genomic analyses of HER3 expression in normal and tumor tissues

All genomic datasets used in our studies are publicly available. The R2 Genomics Analysis and Visualization Platform (http://r2.amc.nl) was used to access the Roth database of normal tissues, TCGA-1097 tumor breast invasive carcinomas and the Brown-198 database of TNBC samples. The normal database was further filtered to only include normal breast tissue (breast, nipple and breast adipose). HER3 (ERBB3), TFR (TFRC) and GLUT1 (SLC2A1) genes were used to interrogate the Normal Endothelial cell (HUAEC/HUVEC)-Luttun-38 and TNBC metastatic brain tumors-Biernat-71 databases, and all normal R2 databases excluding CNS tissue. The latter databases encompass the following: Adrenal (Various-13); B cell (Comerma-24; Johnsen-38; Jima-12; Kauppinen-38; Nussenzweig-8); Blood (Tompkins-857; Uyhelji-555; Jarvela-96; Sindhi-26;Tangye-14; Yamaguchi-30; Yamaguchi-30 fRMA; Fioretos-16; Novershtern-211; Villani-1244; Villani-1140); Colon (Marra-32; Vivier-4); Developmental (embryonic, Tanavde-136; fetal, Wang-4089; fetal, Wang-5290; fetus, Bianchi-40; HES iPSC, Linnarsson-337; HSC, Ogic-21; Stem cell fetal, Xian-24 huex10p; Stem cell fetal, Xian-24 huex10t; embryogenesis, Yi-18; Stem cells, Linnarsson-1715); Endothelial (Luttun-38); Epithelial (Shelhamer-9); Fallopian tube (Shaw-24); Fibroblasts (Mazda-6); Leukocytes (Clark-108; Clark-114; Matthes-33 fRMA; Matthes-33); Liver (McGilvray-8444; McGilvray-4059); Lymphocytes (Goerdt-20; Franken-440; Franken-352; Franken-440; Franken-352; Lye-154; Lye 154 fRMA); Macrophage (Salazar-44); Mesenchymal (Wezel-15 huex10p; Wezel-15 huex10t); Monocytes (PBM-26); Muscle (Gordon-22; Hofman-121 u133a; Hofman-121 u133b; Asmann-40); Pancreatic (Groop-89; Taneera-63); Placenta (Bammler-12); Platelets (Shaw-154); Skeletal (Stephan-4); Spermatogonia (Brinster-6; Spiess-8); T cell (Wicker-35); Thymus (Ferrando-21). Gene expression data is represented as a log2 transformation.

### Flow cytometry detection of HER3

To detect levels of HER3 surface receptors, human [BT549, MDA-MB-231(+) and MDA-MB-231(-)] and murine (4T1) breast cancer cells were incubated with 10 mL of lifting buffer (0.5 M EDTA in PBS) for 1 h at 37°C with gentle agitation. Detached cells were collected, washed with PBS containing 1% MgCl_2_ and 1% CaCl_2_, pelleted and resuspended in PBS, then separated into 1 tube containing 2 x 10^6^ cells. Cells were fixed with 4% PFA for 10 min, rinsed with PBS (3 x 5 min) and suspended in blocking buffer (1% BSA in PBS) at RT. After 1 h, cells were divided into 2 tubes at 1 x 10^6^ cells each, then rinsed with PBS (2 x 5 min). One tube received the Alexa Fluor-488 conjugated anti-HER3 antibody (FAB3481G; R&D Systems, Minneapolis, MN). Cells were incubated with conjugated antibody at 1:100 for 1 h at RT in the dark. Next, cells were rinsed with PBS (3 x 5 min) and analyzed using a benchtop flow cytometer, Moxi GO (Orflo Technologies, Ketchum, ID).

### Human subjects

De-identified specimens were obtained by informed consent under IRB-approved protocol #29973.

### Processing and plating of patient-derived tissue

Surgical specimens placed immediately in cold sterile DMEM after excision were cut into 2-4 mm pieces before undergoing enzymatic and mechanical dissociation using the gentleMACS™ Octo Dissociator multitissue kit and protocol (Miltenyi Biotec). Resuspended cells were then promptly plated into flasks, multiwell plates and/or chamber slides for indicated treatments.

### Immunocytofluorescence/immunohistofluorescence

Cell lines and dissociated patient-derived cells were fixed and processed for immunocytofluorescence, as previously described.^47^

Tissues harvested from mice and human specimens received from fresh cadaver brains (Tissue for Research Ltd.) were preserved in 10% buffered Formalin. Tissues were then paraffin embedded, sectioned, and mounted onto slides. The slides were deparaffinized by incubating in a dry oven for 1 h and then washing slides in xylene 5 times for 4 min each, followed by sequential rinses in 100%, 95%, 90%, 80%, and 70% ethanol, 2× each for 3 min. The slides were then submerged in water. Epitope retrieval was performed by incubating the slides for 30 min at 37 °C in10 mM Tris base, 1 mM EDTA solution, 0.05% Tween 20, pH 9.0. Peroxidase inactivation was performed using 3.0% H_2_0_2_ for 30 minutes. Slides were blocked in 1% BSA and then stained with indicated antibodies against Claudin-5 (Invitrogen 35-2500 1:50), Ad5 (Abcam ab6982 1:100), HIS-tag (Qiagen 34610 1:100) and HER3 (R&D Systems AF4518 1:400). Brain sections were incubated in True Black lipofuscin autofluorescence quencher (Goldbio cat # TBH-250-1).

Where indicated, TUNEL assays were performed according to the manufacturer’s instructions (Roche). Following treatment, slides were counterstained and mounted with DAPI containing Prolong Antifade (Thermofisher). Images of the tissues were captured using a Leica SPE laser scanning confocal microscope and Molecular Devices ImageXpress Pico where indicated. Images were analyzed using ImageJ.

### Recombinant protein production

Recombinant proteins were produced from the pRSET-A bacterial expression vector, which adds an N-terminal hexa-histidine sequence for metal chelate affinity purification of the fusion protein. To produce the fusion protein in bacteria, 50 mL cultures of E. coli BLR(DE3)pLysS transformed with the indicated pRSET constructs were grown to turbidity at 37°C with vigorous agitation, then expanded into 500 mL cultures that were induced with 0.4 mM IPTG at OD_600_ 0.6-0.8. Incubation continued 3 h after induction, at which point cells were pelleted and resuspended in 5 mL lysis buffer (50 mM NaH_2_PO_4_; 50 mM NaCl; pH 8) containing 0.1% Triton X-100 and 1 mM phenylmethylsulfonyl fluoride (PMSF). After one freeze-thaw cycle, 10 mM MgCl_2_ and 0.01 mg/mL DNase I were added, and lysates were gently agitated at room temperature (RT) until viscosity was reduced. Lysates were then transferred to ice, followed by addition of NaCl and imidazole to 1 M and 10 mM final concentrations, respectively. Lysates were clarified by centrifugation at 4 °C, followed by fast protein liquid chromatography (FPLC) affinity purification programmed with a step gradient of imidazole (ranging from 0–500 mM) in 50 nM NaH_2_PO_4_ and 1M NaCl. Fractions containing eluted protein (~92 kDa) were buffer exchanged by ultrafiltration in storage buffer (20 mM HEPES, pH 7.4, 150 mM NaCl, 10% glycerol).

### Particle assembly

Near-infrared (NIR) particles were generated by combining HPK with an Alexa Fluor^®^ 680 end-labeled single-stranded oligonucleotide (ssODN) with sequence: /5’Alex680N/CGCCTGAGCAACGCGGCGGGCATCCGCAAG (obtained from Integrated DNA Technologies) at 4:1 molar ratio of HPK:ssODN and incubating at RT for at least 20 min in HEPES buffered Saline (HBS). Particles were filtered twice using 100k molecular weight cut-off (mwco) ultrafiltration devices. Filtrates and retentates were assessed for protein concentration using a Bradford assay and ssODN payload was assessed based on 680 fluorescence using a Spectramax M2 spectrophotometer.

To make HerDox particles, non-labeled oligonucleotides containing the sequence above and its reverse complement were annealed to form oligonucleotide duplexes as described previously.^48^ HerDox particles were then generated by combining oligonucleotide duplexes with doxorubicin at a molar ratio of 1:10 dsODN:Dox for 10 min at room temperature (RT) with inversion and agitation in TBS (10 mM Tris-HCl, pH 7.4; 150 mM NaCl) followed by adding HPK at a 4:1 ratio of HPK:dsODN:Dox. The mixtures were subjected to ultrafiltration using 100K molecular-weight-cutoff membranes to isolate assembled particles from unassembled components. The dsODN concentration in particles was quantified by spectrophotometric absorbance at 260 nm after heparin-mediated release of nucleic acids as previously described^44^. Doxorubin concentration was determined using absorbance at 480 nm and Beer’s Law using Extinction Coefficient of Dox = 8030 M-1 cm-1. The protein content was measured by spectrophotometric absorbance using the Bradford method (Bio-Rad dye assay).

### Electron microscopy

Particles were prepared for transmission electron microscopy and imaged, as described previously ^49^, through the services of the Electron Imaging Center for NanoMachines within the California NanoSystems Institute at UCLA.

### Structural modeling and molecular dynamics (MD) simulation

The HPK monomeric and pentameric structures were generated as previously described.^44^

### Receptor binding

To analyze HPK binding to tumor cells, cells were plated at 1 × 10^4^/well in 96-well plates and maintained for 24 h before transfer to cold (on ice) to pre-chill cells before receiving 1.5 μg of HPK per well diluted in chilled complete medium. Ligand blocking was performed by pre-adsorbing HPK with soluble human HER3 peptide (Sino Biological) at 10× molar excess in PBS before adding to cells. Cells were incubated on ice for 1 h to promote binding but not uptake, then thoroughly washed with PBS containing 1% MgCl2 and 1%CaCl2, fixed using 4% paraformaldehyde under non-permeabilizing conditions (to detect cell surface proteins only), and processed for ELISA as described previously ^50^. The plates were then processed for crystal violet staining for normalization according to cell number, as described previously ^51^.

### Subcellular fractionation

Subconfluent (70% confluency) HER3+ MDA-MB-435 tumor cells grown in complete media (DMEM, 10% fetal bovine serum, 1% penicillin and streptomycin) were rinsed with 1 X PBS, serum-starved in Buffer A for 1 hour at 37°C, rinsed with 1X PBS, detached with 2 mM EDTA/PBS, then neutralized with double the volume of 1X PBS++. An aliquot of six million cells were washed with PBS and resuspended in 0.7 mL of Buffer A containing 5 nanomoles of indicated protein (quantified by Bradford Assay), incubated with rocking for 1 hour at 4°C to promote receptor binding but not uptake, followed by transfer to 37°C to promote synchronized cell uptake. At the indicated time points, cells were pelleted (10 min, 5000 rpm, 4°C) and washed in a mild acid buffer (1mL of 1X PBS, pH 6) for 5 minutes to remove remaining cell surface protein. Cell pellets were then rinsed with 1X PBS and processed for subcellular fractionation (Qproteome Cell Compartment Kit, Qiagen; following manufacturer’s protocol). Nuclear fractions were isolated and protein precipitated by incubation in 4x volume of ice-cold acetone for 15 minutes followed by pelleting (10 min, 14,000 rpm, 4°C), removal of supernatants and resuspension in storage buffer (10% glycerol and 5% SDS in dH_2_O). Samples were subject to reducing SDS-PAGE and immunoblotted using antibodies recognizing recombinant protein (anti-NRG1 NRG1abcam ab180808 1:5000 dilution in 5% milk) or Lamin A (Invitrogen MA1-06101, 1:1000 in 3% BSA).

### Intracellular trafficking

Intracellular trafficking of HPK was evaluated following our previously established procedures^47^ with the following modifications: 12-well plates containing 10,000 cells/well plated on coverslips were briefly pre-chilled then exposed to 7 μg HPK per well in Buffer A for 1 h to promote receptor binding but not internalization while equivalent samples received 100 nM bafilomycin-A1 in Buffer A for 30 min before adding HPK. Plates were then transferred to 37°C to promote synchronized uptake and intracellular trafficking. Cells were fixed at indicated time points after warming and then processed for immuno-identification of HPK using an antibody that recognizes the polyhistidine tag (RGS-His antibody; Qiagen 1:100) and counterstained for nuclei using DAPI. Images were acquired using a high-throughput digital microscope (Molecular Devices ImageXpress^®^ Pico Automated Cell Imaging System) using a 40X magnification lens. Exposure times for each fluorescence wavelength remained fixed to compare between treatments and timepoints.

Endosome maturation staining antibodies for RAB7 and EEA1 were purchased from Abcam (ab50533 and ab206860, respectively). Samples were imaged using a Leica SPE laser scanning confocal microscope. Acquired images were imported and split into individual channels. Individual cells in selected channels were delineated and pixel overlap was evaluated using ImageJ.

### Isothermal titration calorimetry (ITC)

The ITC experiments were performed using a MicroCal PEAQ-ITC at 25°C. For measuring heat changes during assembly of HPK with ssODN, the ssODN was suspended in RNA annealing buffer (10 mM Tris-Cl, pH. 7.5; 50 mM NaCl, and 1 mM EDTA all in ultrapure nuclease free water.).

To avoid buffer mismatch, HPK was buffer exchanged into the same RNA annealing buffer. The ssODN (300 μL of 1.0 μM concentration) was placed in the stir cell and titrated with 5.4 μM HPK at 2 μL HPK/titration in a 2.5 min injection for the titration peak to return to the baseline. The K_d_ was calculated using the MicroCal PEAQ-ITC analysis software, as well as Prism GraphPad software, using the one-site model. Control experiments included titrating HPK into buffer, buffer into the ssODN, and buffer into buffer to ensure that any measured changes were not due to buffer mismatches or other artefacts. The three controls were used in a composite to subtract the heat of dilution and background noise from the baseline. ITC measurements of HPK assembly with dsODN and of HerDox assembly are described in the **Supplemental Methods**.

### Dynamic light scattering (DLS)

A Malvern ZEN 3600 Zetasizer Nano was used for DLS analyses. A typical analysis comprised three or more measurements per sample, each measurement comprising 100 runs, with an average of 34K particle counts/sec (kpcs). The reported average is the number particle size determination parameter, which yields the most frequent particle size in the sample accounting for intensity fluctuations of larger particles. The intensity of the particles was computed via Zetasizer Software version 7.01, which applies the Stokes-Einstein equation to correlate the change in the scattering intensity and particle movements.

### Serum digest (protection) assay

Free dsODN alone (60 pmol) or preincubated with HPK (2 μg for 30 min at RT) was incubated in whole (100%) active mouse serum at 37°C for 1 h and then subject to agarose gel electrophoresis.

### BBB chip

The chip is composed of a flexible polydimethylsiloxane (PDMS) elastomer that contains two closely opposed and parallel micro-channels (1 × 1 mm and 1 × 0.2 mm, brain and blood channel, respectively)^52^ separated by a laminin-coated, porous flexible PDMS membrane (50 μm thick, with 7-μm diameter pores with 40 μm spacing, resulting in 2% porosity over a surface area of 0.171cm2 separating the two channels). Cell aggregates (“EZ-spheres”)^53^ were derived from cells obtained through IRB # 21505. EZ-spheres were dissociated into single cells using accutase and were seeded into BBB chips at a density of 1.25E6 cells/mL in TDM (terminal differentiation media containing Rock inhibitor 1:2000 (Stemgent). Cells were allowed to settle for 2hrs, and then flushed with TDM without Rock inhibitor. Media was replaced with 100uL of TDM every other day. Five days later, iBMECs (brain microvascular endothelial cells) were seeded into the bottom channel. iBMECs were seeded at 15E6 cells/mL in S3 BMEC media containing Rock inhibitor (1:2000) and inverted for 2hrs. A second seeding was performed after the two hours with the same protocol but allowed to sit upright for 2hrs. Following the second incubation period, the chips were flushed with S3 BMEC media without Rock inhibitor. The following day, chips were flushed with fresh TDM and S4 BMEC media. On the next day, chips were added to Emulate, Inc. pods and placed on an active flow of 30uL/Hr. Chips were first validated via paracellular permeability assays using dextran-FITC overnight to determine barrier function of the chips under flow. Validated BBB chips were treated with NNCs at a concentration equating 1μg/mL of HPK that was passed through the endothelial channel with or without 10:1 blocking peptide commercially purchased from Sinobiological (10201-H08H). After 4 hours of constant flow, the chips were fixed using 4% PFA and subjected to immunocytofluorescent staining.

### Sandwich ELISA

Non-tissue culture treated polystyrene plates were pre-coated using Rabbit Anti-Ad5 (Abcam-ab6982) diluted in pH 9.2 Sodium Bicarbonate. Plates were then blocked using 3% Bovine Serum Albumin (BSA). After washing with PBS 3 times, BBB effluents were incubated overnight. Signal was detected using mouse anti-His antibody from Qiagen (Cat No./ID: 34660 1:500). Anti-mouse HRP was used for colorimetric development (Cat No./ID-A8924 1:1000). A650 reads were started 15 minutes after addition of TMB. Once developed and read using Molecular Devices SpectraMAx M2, the reaction was stopped by adding 1N HCl and the A450 was acquired. An HPK standard curve was processed in parallel for quantifying HPK NNCs collected from BBB chip effluents.

### Animal subjects

Immunodeficient (NU/NU) and immunocompetent (BALB/c) mice were obtained from Charles River Laboratories, Inc. All procedures involving mice were performed following IACUC-approved protocols #6037 and #5790 in accordance with the institutional and national guide for the care and use of laboratory animals. The criteria for euthanasia include tumor ulceration, interference with ambulation and access to food and water, or exhibit a Body Condition Score (BCS) of less than 2 (emaciation, prominent skeletal structure, little/no flesh cover, visible and distinctly segmented vertebrae).^54–55^

### Tumor models

Peripheral breast tumor models used for biodistribution studies were established in 6-week-old female mice. For xenograft models, immunodeficient (Nu/Nu) mice received bilateral flank implants of JIMT-1 human HER2+ tumor cells (1e7 cells/implant). For immune-competent models bearing peripheral TNBC tumors, BALB/c mice received bilateral mammary fat pad injections of 4T1-LucGFP cells (1e4 cells/injection in 0.1 mL PBS). Tumor volumes (height x width x depth) were monitored ~3x/week under single-blinded conditions (treatment groups unknown to the individual acquiring measurements). Mice were randomized at tumor establishment (≥100–150mm3) into separate treatment groups (n=5 mice per group).

For mice receiving intracranial tumors, anesthetized 4-week old female Nu/Nu mice were positioned using a stereotactic frame to drill a burr hole in the skull using a steel bit positioned 2 mm right of the sagittal and 2 mm anterior to the lambdoid suture. Ten-thousand 4T1-LucGFP cells were then implanted using a stereotactic frame to guide a Hamilton syringe. After implantation, bone wax was used to seal the orifice and a surgical staple used to seal the incision, which could be removed 5 days later. Treatments and group size were as follows: n=12 for HerDox and Lipodox, n=14 for Mock. Tumor growth was monitored via luciferase beginning on day 4 after implantation and mice randomized before treatments.

### Luciferase Imaging

Luciferase monitoring was performed every four days beginning on day 4 post-implantation. Mice received 200 uL I.P. of 30 mg/mL D-Luciferin (Caliper) dissolved in PBS 10 minutes before imaging using IVIS. D-Luciferin was allowed to circulate in the animals for 15 minutes followed by imaging using a Perkin-Elmer IVIS Lumina Spectrum. Signal intensities were quantified using region of interest tools in LivingImage™ software.

### MRI

A 9.4T MRI (BioSpec 94/20USR, Bruker GmhB) was used for imaging tumor locations and volumes. The tumor perimeters were visually aided by i.v. injection of Gadovist contrast agent, 7.5 μmol. Mice were imaged under inhaled isofluorane anesthetic, 1.7%. Images were collected with an in-plane resolution of 70 μm using an acquisition matrix of 256 x 196 and zero filling in the phase encoding direction to 256, FOV of 1.80 cm x 1.80 cm. Twenty consecutive 0.7 mm slices covered the tumor implanted region. Two averages were collected with a repetition time of 750 ms and an echo time of 8.77 ms for a total scan time of 4.9 minutes using a mouse 4-channel brain array coil (T11071V3, Bruker GmbH) for reception and a whole-body transmission coil (T10325V3, Bruker GmbH) for excitation. Volume calculations for positive contrast brain regions were determined by integration over the entire tumor by a blinded analyzer. For each slice containing tumor enhanced regions, a region of interest (ROI) was drawn to encompass the area. The area of each slice was multiplied by the slice thickness and the individual volumes were summed to determine the area of the tumor. Analysis utilized the Bruker Paravision 5.1 software. MRI core staff performed quantifications in a blinded fashion.

### Biodistribution

For quantifying particle delivery by ICP-MS, each mouse was administered a single tail vein injection of HPK bioparticles loaded with a gallium(III) metallated corrole (HPK-S2Ga or HerGa)^42, 49^ or S2Ga alone at 1.5 nmoles S2Ga per injection. At predetermined time periods (6, 12, and 24 h), (n=2 per treatment) mice were euthanized and the major organs (brain, heart, kidneys, liver, lungs, spleen, and tumors) were excised, weighed and transferred to the University of California Los Angeles ICP-MS core facility where samples were digested overnight and processed by ICP-MS to measure the tissue content of Ga(III) metal.

For mice receiving NNCs (n=5), each mouse was administered a single tail vein injection of HPK bioparticles loaded with near-infrared (Alexa Fluor 680) -labeled oligonucleotides at a dose equating 1.50 nmoles oligonucleotide/injection. Mice were monitored by epifluorescence imaging at the indicated time points after injection using a IVIS^®^ Spectrum In Vivo Imaging System (PerkinElmer, Waltham, MA), followed by tissue harvest and acquisition of average radiant efficiency per tissue. Where indicated, relative tissue content of NIR-ODN was determined based on extrapolating measurements acquired from extracted tissue against a standard curve.

For mice (n=5 each treatment) receiving directly labeled protein or particles (NIR-labeled HPK capsomeres, Tz, or BSA), each protein was delivered by single tail-vein injection at 12 nmol of labeled protein/injection. Proteins were labeled with Alexa Fluor-680 at primary amines and isolated from unconjugated dye by size exclusion chromatography using a commercial protein labeling kit, following the manufacturer’s protocol (LifeTechnologies).

### Statistical methods

Except where indicated, in vitro data are presented as the mean of triplicate samples ± standard deviation from at least three independent experiments. For normally distributed in vitro data, significances were determined by one-way ANOVA followed by Tukey post hoc analyses unless otherwise indicated. In vivo data are presented as normalized mean ± standard deviation. Non-parametric analyses were used to determine significances within *in vivo* experiments where appropriate using Kruskall-Wallis tests followed by Mann-Whitney post hoc analysis.

## RESULTS

### HPK bioparticles containing oligonucleotide cargo form HER3-targeting nano-nucleocapsids (NNCs)

Bioinformatic and clinical sources support the evidence for HER3 as a potential homing beacon for targeting metastatic and triple-negative breast tumors. Tumor cases classified as invasive breast carcinoma (N=1,100) in The Cancer Genome Atlas (TCGA) databases exhibited significantly increased HER3 expression compared to normal breast tissue (**Fig. 1A**). Additionally, 198 cases from the same database exhibited a triple-negative phenotype of below average estrogen receptor (ER), progesterone receptor (PR) and HER2, of which 81% of cases showed above average HER3 expression (**Fig. 1B**). In agreement with these clinical findings, human TNBC lines BT549 and MDA-MB-231 (MDA-231+) and the mouse TNBC line 4T1 display substantial levels of cell surface HER3 (**Fig. 1C**) in contrast to an MDA-MB-231 sub-line (MDA-231-) that lacks detectable HER3 (**Fig. 1, C-D**). Patient-derived HER2+ breast tumor cells escaping HER2-targeted intervention also displayed increased HER3 compared to normal breast cells obtained from the same patient (**Fig. 1E**).

**Figure 1.**
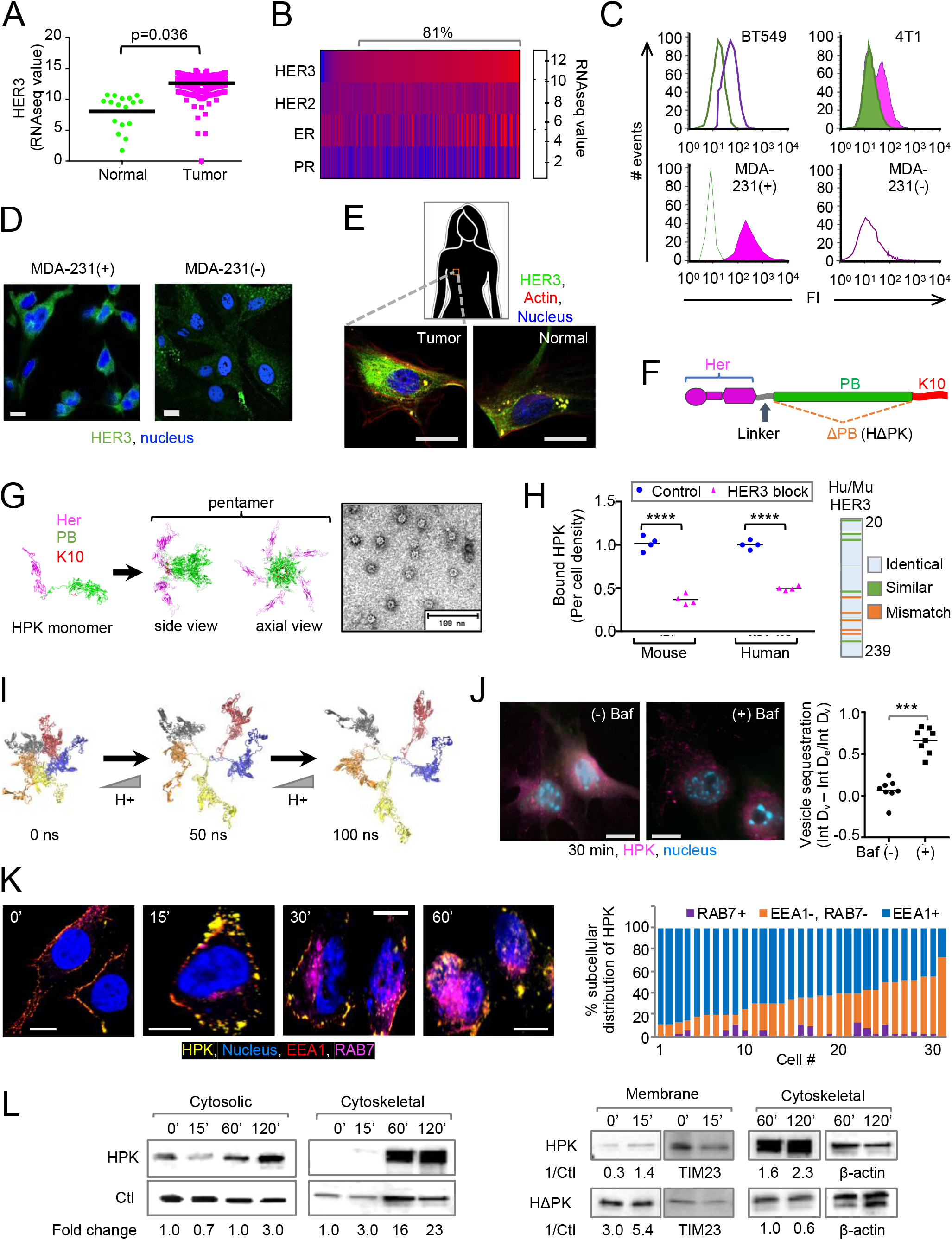
HER3 expression and HPK penetration of invasive breast tumor cells. **A,** Bioinformatics analysis comparing HER3 gene expression of all invasive breast cancers including TNBC (TCGA-1097 database; N=199 samples) against normal breast tissue (Roth database; N=17 samples) using the R2 Genomics Analysis and Visualization Platform (http://r2.amc.nl). **B,** Gene expression heatmap of 198 TNBC tumors (Brown database) queried for HER3 (ERBB3), HER2 (ERBB2), estrogen receptor (ESR1) and progesterone receptor (PR) expression. Samples are ranked by HER3 expression levels. Percent of cases showing above average HER3 (defined by average HER3 expression in normal breast) are demarcated above the heatmap. **C,** Flow cytometry measurement of cell surface HER3 on human [BT549, MDA-231(+), MDA-231(-)] and mouse (4T1) TNBC lines. **D,** Immunocytofluorescence of HER3+ TNBC cells [MDA-231(+)] in comparison to respective cells with no/low HER3 expression [MDA-231(-)]. Scale bar, ~8 μm. **E,** Immunocytofluorescence of patient-derived tumor cells. Scale bar, ~8 μm. **F,** Graphical representation of the HPK linear sequence from amino [N] to carboxy [C] terminus (left to right), highlighting: HER3-binding motif (Her) appended to a flexible linker sequence comprised of neutral residues (Gly-Gly-Ser)_2_ followed by the penton base sequence (PB) and decalysine (K10). **G,** Ribbon models of monomeric and pentameric HPK generated by molecular dynamics (MD) simulation, with each functional region depicted by a designated color. Inset, transmission electron micrograph (TEM) of HPK capsomeres. **H,** Cell surface ELISA (graph) showing HPK binding to human (BT549) and mouse (4T1) TNBC cells -/+ competing HER3 peptide. ****, p<0.0001. Data show individual measurements from quadruplicate samples and corresponding means. Inset, graphical alignment of mouse and human HER3 ligand binding domains (amino acid residues 8-239) showing identical, similar, and mismatched residues. **I,** Video stills of MD-simulated HPK pentamer (with each monomer distinguished by a different colored ribbon structure) in a neutral solution with titrating H+ added over time. Full video of capsomere dynamics is shown in ***Supplemental Movie 1*. J,** Immunocytofluorescence of 4T1 cells at 30 min after uptake of HPK capsomeres -/+ bafilomycin-A1. Scale bar, ~8 μm. Graph summarizes contrast between vesicular (v) and extravesicular (e) regions by measuring integrated densities (Int D) of each and applying the formula shown by the y-axis label. **K**, Intracellular trafficking of HPK capsomeres in HER3+ MDA-MB-435 cells in relation to early endosomes (EEA1) and late (RAB7) endo-lysosomes. Scale bar, ~5 μm. Graph summarizes the intracellular distribution of HPK per cell across all time points as a % of internalized HPK. **L,** Immunoblots of subcellular fractions isolated from HER3+ MDA-MB-435 human tumor cells harvested and processed at the indicated time points during uptake of HPK or HΔPK. Relative levels of uptake are quantified by normalizing band densitometries with those of respective fraction controls. Where indicated, fold change is reflected by the difference in normalized band densitometry at subsequent time points compared to that at 0 min.

The recombinant multidomain protein, HPK, contains the receptor-binding region of the HER3 ligand, neuregulin-1α (amino acids 35-239, comprising the Ig-like and EGF-like domains)^56^, expressed as an amino- [N] terminal fusion to a membrane-penetrating moiety derived from the adenovirus penton base capsid protein which in turn is modified by a carboxy- [C] terminal decalysine tail (**Fig. 1F**) ^44^. The penton base domain of HPK drives the formation of highly stable pentameric barrels appearing as donut or ring -like structures under TEM (**Fig. 1G**) reminiscent of viral capsomeres that form naturally upon protein expression ^57^. HER3 specificity and species cross-reactivity is confirmed by the ability of a human HER3 peptide to block binding of capsomeres to both mouse and human HER3-expressing TNBC cell lines (**Fig. 1H**). This is consistent with the high amino acid sequence identity (94%, within extracellular domains I and II)^58^ shared by both human and mouse HER3 (**Fig. 1H, inset**).

The endosomolytic mechanism of HPK is thought to occur in part by protonation of solvent-accessible residues lining the lumen of the capsomere barrel when exposed to the decreasing pH of the endosome,^44^ causing charge-mediated dispersion of constituent monomers and exposure of hydrophobic domains mediating pentamerization. Charge-mediated dispersion is predicted by molecular dynamics (MD) simulation of HPK pentamers in solution which shows that a decrease in pH disrupts pentamerization and disperses the monomers from one another thus exposing intra-pentamerization domains (**Fig. 1I** and **Supplemental Movie 1)**. Exposure of these hydrophobic domains would allow interactions with endosomal membrane lipids leading to membrane destabilization under acidic pH. Consistent with this, the proton pump inhibitor bafilomycin prevents the diffusion of HPK into the cytoplasm of 4T1 TNBC cells while retaining HPK sequestration in vesicle-like structures suggesting that endosomal penetration is pH-dependent (**Fig. 1J**). This is consistent with the intracellular trafficking profile of HPK, which avoids overlap with the lysosomal biomarker, Rab7, while entering early (**Fig. 1K**) and post-endosomal fractions including the cytosol and cytoskeleton (**Fig. 1L** and **Supplemental Fig. S1**).^44^ Post-endosomal trafficking of HPK agrees with the trafficking route of soluble penton base proteins which transit along microtubules toward the nucleus after endosomal escape.^44, 47, 59^ Here we show that in contrast to HPK a penton base-deleted construct HΔPK (**Fig. 1F**) shows early sequestration in the membrane compartment and considerably reduced arrival at late (cytoskeletal) trafficking compartments (**Fig. 1L** and **Supplemental Fig. S1**), indicating that post-endosomal delivery requires the penton base domain.

Computational modeling has shown that the formation of HPK capsomeres aligns the decalysines on one end of the barrel^44^ creating a positively-charged surface that repels the capsomeres from one another in solution (**Fig. 2A**). Exposure of HPK capsomeres to anionic cargo such as small nucleic acids neutralizes the charged surfaces of the barrels thus allowing capsomeres to converge around the cargo (**Fig. 2A**).^44^ This occurs at a nearly 1:1 spontaneous binding ratio in the nanomolar range when HPK is mixed with oligodeoxynucleotide (ODN) cargo (**Fig. 2B**). The shape complementarity of the capsomeres and interaction between inter-capsomere binding motifs (highlighted in **Supplemental Fig. S2A**) drives their assembly into polyhedra (as seen under TEM) containing the encapsulated cargo (**Fig. 2A**). This is supported by the observation that HPK in solution shifts to an increase in hydrodynamic diameter upon exposure to ODN (**Fig. 2C** and **Supplemental Fig. S2B**) in agreement with the aforementioned TEM (**Fig. 2A**) and is validated by the ability of HPK to retain fluorescently tagged ODN on ultrafiltration membranes (**Fig.2D**). The resulting bioparticle assembly protects encapsulated nucleic acid cargo from serum endonuclease digestion (**Fig. 2E**) and creates a capsid-like morphology distinguishing these bioparticles as “**nano-nucleocapsids**” (**NNC**s). The predicted display of HER3 ligands distributed over the particle surface (**Fig. 2A**) is supported by the accumulation of HPK NNCs at HER3 sites when exposed to primary breast tumor cells freshly excised from a HER2+ patient (**Fig. 2F**).

**Figure 2.**
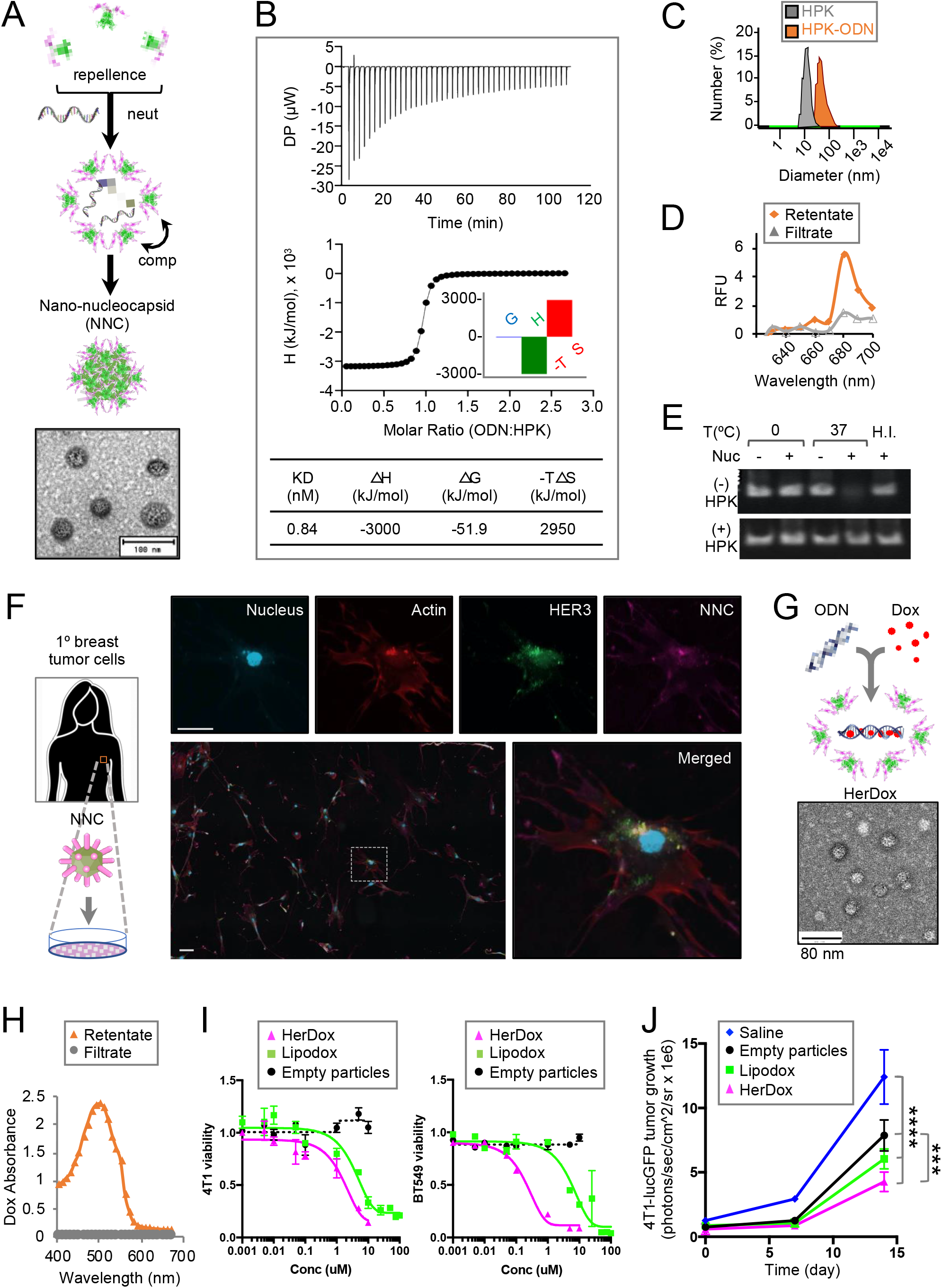
Formation of HER3-targeting nano-nucleocapsids (NNCs). **A,** Graphical representation of NNC assembly (showing molecular dynamics-generated capsomeres) nucleated by the cargo-driven convergence of capsomeres. Inset, transmission electron microscopy (TEM) of NNCs. **B,** Isothermal titration calorimetry of binding interaction between HPK and single-stranded oligonucleotide (ssODN). Top panel shows the raw thermogram generating the binding isotherm (lower panel) and signature plot (inset) summarizing the thermodynamics parameters of the reaction (inserted table). **C,** Hydrodynamic diameters of NNCs containing ssODN (HPK-ODN) in comparison to HPK alone, as determined by dynamic light scattering. **D,** Fluorimetry of NNCs containing NIR-labeled oligodeoxynucleotide (NIR-ODN) after filtration to isolate assembled particles (retentate) from unassembled NIR-ODN (filtrate). **E,** Gel electrophoresis and ethidium bromide staining of dsODN after incubation in serum containing active nucleases (nuc) -/+ pre-assembly with HPK. H.I., heat-inactivated serum. **F,** Binding of HPK particles to patient derived tumor cells. Breast tumor cells were isolated from dissociated tissue freshly acquired from HER2+ patient. Delineated region is enlarged in top row of panels and lower right panel. Scale bar, ~25 μm. **G,** Graphical representation of HerDox assembly and TEM of HerDox (inset). **H,** Spectrophotometric absorbance of HerDox after filtration to isolated assembled particles (retentate) from unassembled Dox (filtrate). **I,** HerDox killing curves on mouse (4T1) and human (BT549) TNBC lines comparing HerDox to Lipodox and empty (no Dox) NNCs. Cytotoxicity was measured by crystal violet staining on cells that were fixed at 24 h after addition of indicated reagents, as previously described.^49^ Data represent mean±SD of 3 independent experiments performed in triplicates. **J,** Growth of subcutaneous bilateral 4T1 tumors (determined by acquisition of tumor bioluminescence) in BALB/c mice during systemic treatment with indicated reagents (regimen described in ***Supplemental Methods***). Day 0 corresponds to first day of treatment (~5 days after tumor implant). ***, p=0.0005. ****, p<0.0001. Data represent mean±SD of 5 mice per treatment group.

Intercalating the ODN with doxorubicin (Dox) (reaction shown in **Supplemental Fig. S2C**) before assembly with HPK (reaction shown in **Supplemental Fig. S2D**) forms drug-loaded NNCs designated HerDox that retain a similar polyhedral shape and size as drug-lacking NNCs (**Fig. 2G** and **Supplemental Fig. S2E**).^42^ The assembly reaction occurs at a 6:1:7 molar stoichiometry of HPK:ODN:Dox in the nanomolar range (**Supplemental Fig. S2D**) with assembled particles showing no detectable leakage of the drug during ultrafiltration (**Fig. 2H**). Therapeutic efficacy of HerDox compared favorably against liposomal doxorubicin (Lipodox) in reducing growth of HER3+ TNBC cells in culture (**Fig. 2I**) and in reducing growth of peripheral TNBC tumors *in vivo* during systemic delivery (**Fig. 2J** and **Supplemental Fig. S3**) while empty particles (NNCs lacking the drug) showed no growth-promoting effect (**Fig. 2, I-J** and **Supplemental Fig. S3**).

### Systemic NNCs enter mouse brain parenchyma

We systemically administered NNCs carrying near-infrared (NIR)-labeled ODN in mice bearing peripheral HER2+/HER3+ JIMT1 tumors to track their tissue distribution. Systemic NNCs not only exhibited tumor-preferential accumulation, but also coincided with the brain in contrast to the negative control molecule, trastuzumab (Tz) (**Fig. 3A**), which homes to HER2+ tumors but does not cross the BBB.^60–61^ These findings were echoed by HPK capsomeres directly labeled with a NIR dye which could be detected in the brains of tumor-bearing (**Fig. 3B**) and tumor-free (**Supplemental Fig. S4, A-B**) mice in contrast to labeled Tz (**Fig. 3, A-B** and **Supplemental Fig. S4, A-B**) and non-targeted protein (BSA) (**Supplemental Fig. S4, A-B**). These findings were also validated using inductively coupled plasma mass spectrometry (ICP-MS) to measure tissue content of HPK bioparticles delivering a metallated probe (gallium[III] corrole or S2Ga)^42^ in a syngeneic mouse model of TNBC (**Fig. 3C**). In contrast to the untargeted probe alone, delivery by HPK resulted in accumulation of the Ga(III) in mouse brain tissue as well as peripheral tumors (**Fig. 3C**).

**Figure 3.**
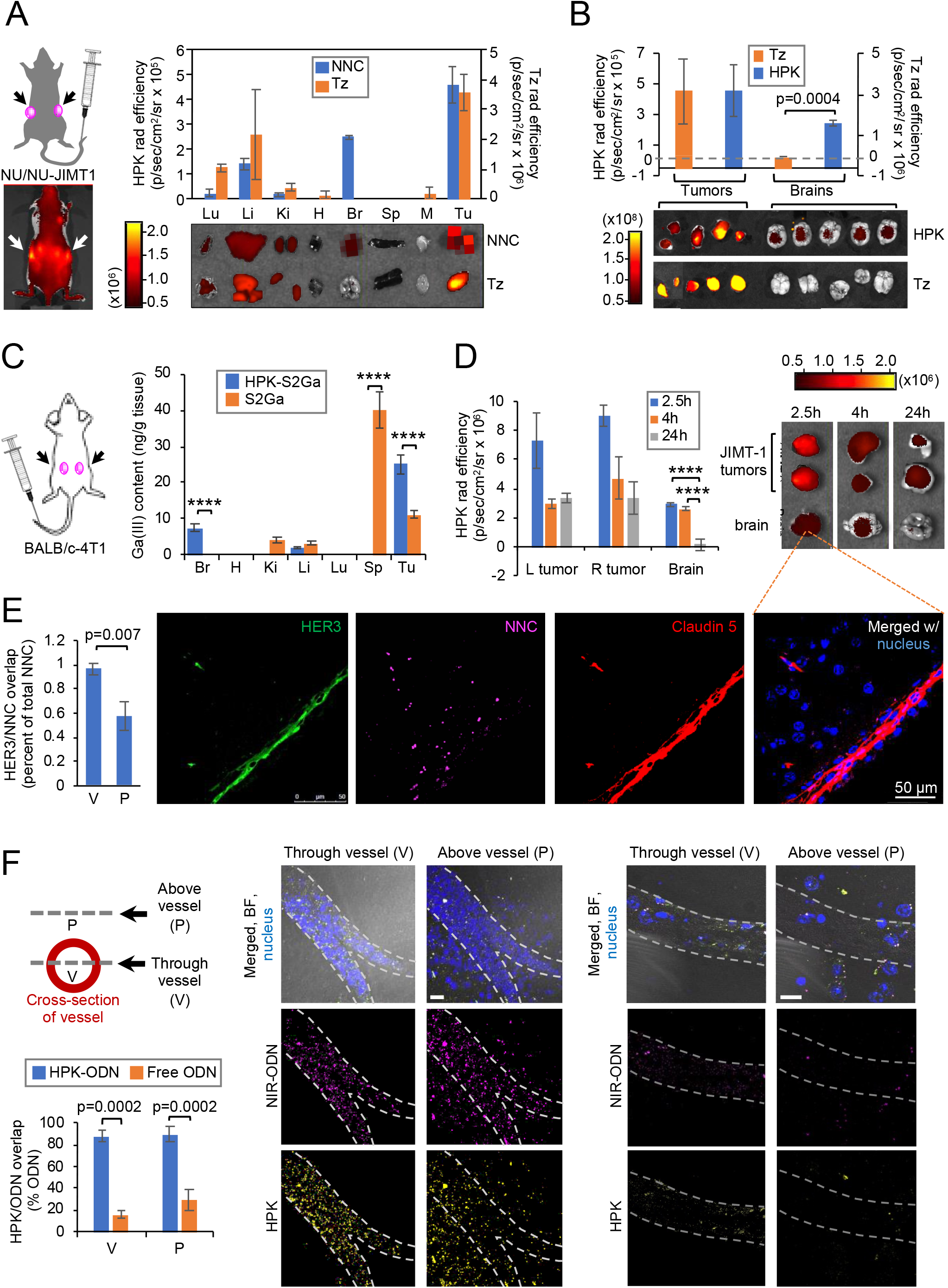
Systemic NNCs enter mouse brain parenchyma. **A,** Tissue distribution of near infrared (NIR)-labeled NNCs or trastuzumab (Tz) after systemic delivery in mice bearing subcutaneous JIMT-1 breast tumors. Arrows point to tumor locations in inset. Graph summarizes average radiant efficiencies collected from tissue harvested >4h after systemic delivery of each reagent. Data represent mean±SD of each corresponding tissue from 5 mice per treatment group. Tissues shown below the x-axis were acquired from representative mice of each cohort. Lu, lung; Li, liver; Ki, kidney; H, heart; Br, brain; Sp, spleen; M, muscle; Tu, tumor. **B,** Brain and tumor distribution of near infrared (NIR)-labeled HPK or trastuzumab (Tz) after systemic delivery in mice bearing subcutaneous JIMT-1 breast tumors. Graph summarizes average radiant efficiencies collected from tissue (shown below x-axis) harvested 2.5 hours after systemic delivery of each reagent. Data represent mean±SD of each corresponding tissue from 5 mice per treatment group. **C,** Tissue content of HPK bioparticles after systemic delivery of a gallium(III)-metallated corrole payload (S2Ga) in an orthotopic syngeneic mouse model of TNBC. Inductively coupled plasma mass spectrometry (ICP-MS) was used to measure Ga(III) content from each digested organ or tumor. Data represents 3 separate measurements of each organ harvested from two mice each per treatment. ****, p<0.0001. **D,** Quantification of tumor and brain distribution at indicated time points after systemic administration of NNCs delivering NIR-ODN in mice bearing subcutaneous JIMT1 tumors. ****, p<0.0001. Data represent mean±SD of 12 samples at each time point. Images show tumors and brains from representative mice at each time point. **E,** Imaging and quantification of NNC and HER3 localization in relation to brain vasculature (delineated by claudin 5) at 2.5h after systemic delivery of NNCs delivering NIR-ODN. Images show channel separated and merged micrographs capturing representative mouse brain section after immunohistofluorescent stain of indicated biomarkers. Graph shows quantification of NNC and HER3 overlap in brain vessel (V) compared to parenchyma (P) from 3 independent specimens. Data represent mean±SD. **F,** Assessment of HPK and NIR-ODN overlap at the vasculature of two representative brain sections from experiment in ***E*** showing adjacent z-axis planes that transect the vessel (V) and the parenchyma above the vessel (P). “Free ODN” designates the detection of NIR-ODN without overlapping HPK. Graph summarizes the overlap of HPK with NIR-ODN at each location and shows the mean±SEM from 3 independent specimens. Scale bar, ~20 μm.

In mice bearing peripheral JIMT1 tumors, systemic NNCs were detected in tumors and brains at 2.5 hours and 4 hours after administration and remained detectable in tumors but not in brains by 24 hours (**Fig. 3D**). Likewise, systemic NIR-labeled HPK capsomeres remained detectable in brains of tumor-free immunocompetent mice up to 4h after administration and cleared from brains by 24h (**Supplemental Fig. S4C**). Immunohistologic evaluation of brains collected at 2.5 hours after systemic administration showed that HER3 is present at robust levels on the brain vasculature (delineated by the tight junction marker claudin-5) and that the NNCs coincided with both the vasculature and extravascular parenchyma (**Fig. 3E**). To more precisely assess the locality of HPK NNCs and their retention of cargo with respect to the brain vasculature, we analyzed adjacent visual planes along the z-axis of the microscopic viewing depth focusing on the plane transecting the vessel (V) and that above the vessel (P) in the extravascular parenchyma (**Fig. 3F, schematic**). Detection of both HPK and fluorescently-tagged cargo (near-infrared labeled oligonucleotide, or NIR-ODN) within and above the vessel (**Fig. 3F, micrographs**) enabled assessment of HPK/cargo overlap or non-overlap at each spatial location. HPK and NIR-ODN coincided both within the vessel (**Fig. 3F, graph, “V”**) and within the parenchyma (**Fig. 3F, graph, “P”**) while non-overlapped NIR-ODN was low to negligible in both locations.

### HER3 mediates passage across the human BBB

These findings led us to examine whether HER3 is typically present on normal brain endothelium. We found that HER3 is prominently expressed on both healthy (non-tumor) adult mouse and human brain vasculature (delineated by claudin 5) whereas HER3 is nearly undetectable in the extravascular brain parenchyma of both species (**Fig. 4, A-B**). Human gene expression analyses show that HER3 compares favorably against conventional BBB transporter genes encoding transferrin (TfR) and glucose (GLUT1) receptors for targeting to brain-localized tumors. Specifically, HER3 is significantly lower than TfR and GLUT1 on peripheral endothelia (**Fig. 4C**) while showing significant increase over TfR and GLUT1 on brain metastatic breast tumors (**Fig. 4D**). Normal non-CNS tissue also shows an overall HER3 expression that is significantly lower than TfR and GLUT1 (**Fig. 4, D-E**).

**Figure 4.**
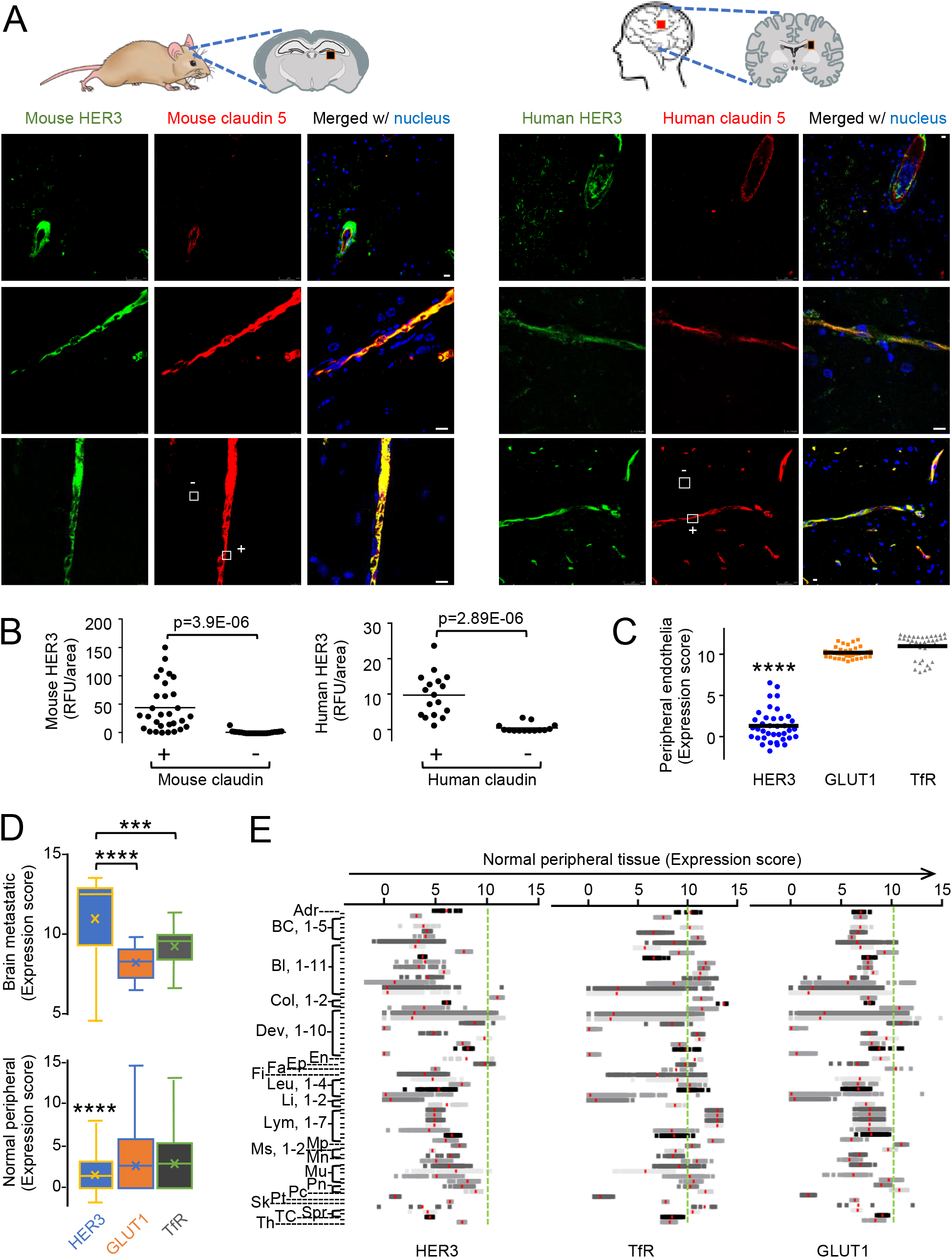
HER3 expression on the blood brain barrier (BBB). **A,** Immunohistofluorescence of frontal cortex from non-diseased adult murine and human brains showing cross sections and longitudinal views of blood vessels within brain specimens. Specimens were obtained from female adult immunodeficient mice (6+ months) and human male frontal cortex, ages 71, 68, and 76 years old (Tissue for Research Ltd.). White squares delineate claudin positive (+) and negative (-) areas. Scale bar, ~10 μm. **B,** Quantification of overlap between claudin-5 and HER3 in specimens shown in ***A***. Data points represent the means of individual regions of interest in each indicated zone. **C,** R2 database analysis of HER3 (N=38), glucose receptor (GLUT1; N=38) and transferrin receptor (TfR; N=38) gene expression in peripheral (non-brain) endothelial tissue, showing mean and individual data points. ****, p<0.0001. **D,** R2 database analysis of HER3, GLUT1, and TfR gene expression in human breast cancer brain metastases (upper graph) and human peripheral (non-brain) non-tumor tissues (lower graph, summarizing collective expression scores shown in ***E***). Data represent mean (X) with median and interquartile range. ***, p=0.0006. ****, p<0.0001. N=31,408 per group. **E,** Itemized expression scores of human peripheral (non-brain) non-tumor human tissues comparing HER3, TfR and GLUT1 genes. Red line in each category indicates the mean. Dashed vertical line delineates threshold for high expression. Each y-axis tick mark represents a separate database. Multiple databases within the same category are enumerated. Sources and N of each database are listed in the *Methods*. Adr, adrenal; BC, B cell; Bl, blood; Col, colon; Dev, developmental; En, endothelial; Ep, epithelial; Fa, fallopian tube; Fi, fibroblasts; Leu, leukocytes; Li, liver; Lym, lymphocytes; Mp, macrophage; Ms, mesenchymal; Mn, monocytes; Mu, muscle; Pn, pancreatic; Pc, placenta; Pt, platelets; Sk, skeletal; Spr, spermatogonia; TC, T cells; Th, thymus.

To examine whether NNCs delivering NIR-ODN extravasate in human tissue, we used a reconstituted human BBB (BBB-on-a-chip) derived from induced pluripotent stem cells (iPSCs) differentiated into apposing neuronal and endothelial layers recapitulating the brain-vascular interface.^62^ Channels creating the closed vessel and overlaying neuronal tissue enable flow of molecules through the endothelial tube (**Fig. 5A**), whose integrity is validated using fluorescently-labeled dextrans of varied sizes.^62^ Consistent with the histologies of both mouse and human endothelia observed earlier, the human BBB-chip displayed considerable levels of HER3 on the endothelial surface (E) in contrast to the neuronal layer (N) (**Fig. 5B**), and HER3 was significantly higher on the endothelial surface abutting the neuronal layer (proximal) in contrast to the distal endothelial surface (**Fig. 5, B and C**). HPK NNCs flowing through the endothelial tube showed considerable localization at the endothelial-neuronal (proximal) interface in contrast to the distal endothelial surface, concomitant with the contrasting HER3 levels on these surfaces (**Fig. 5, B and D**). Approximately 42% of the NNCs injected into the endothelial tube could be collected from the neuronal chamber effluent (**Fig. 5, B and D**) suggesting that NNCs can transit the endothelial barrier. A HER3-blocking peptide significantly reduced the emergence of NNCs into the neuronal layer and reduced NNC accumulation in the neuronal chamber effluent (**Fig. 5, E-F**) suggesting that BBB transit is mediated by HER3. As nucleic acids are unable to cross the vasculature or penetrate into cells on their own, a free NIR-ODN control enabled us to further confirm the integrity of the endothelial tube and assess whether HPK can enable passage of NIR-ODN across the endothelial barrier. Indeed we found that NIR-ODN alone remained largely retained in the endothelial tube whereas the NIR-ODN emerged into the neuronal (N) layer when delivered by HPK NNCs (**Fig. 5G**).

**Figure 5.**
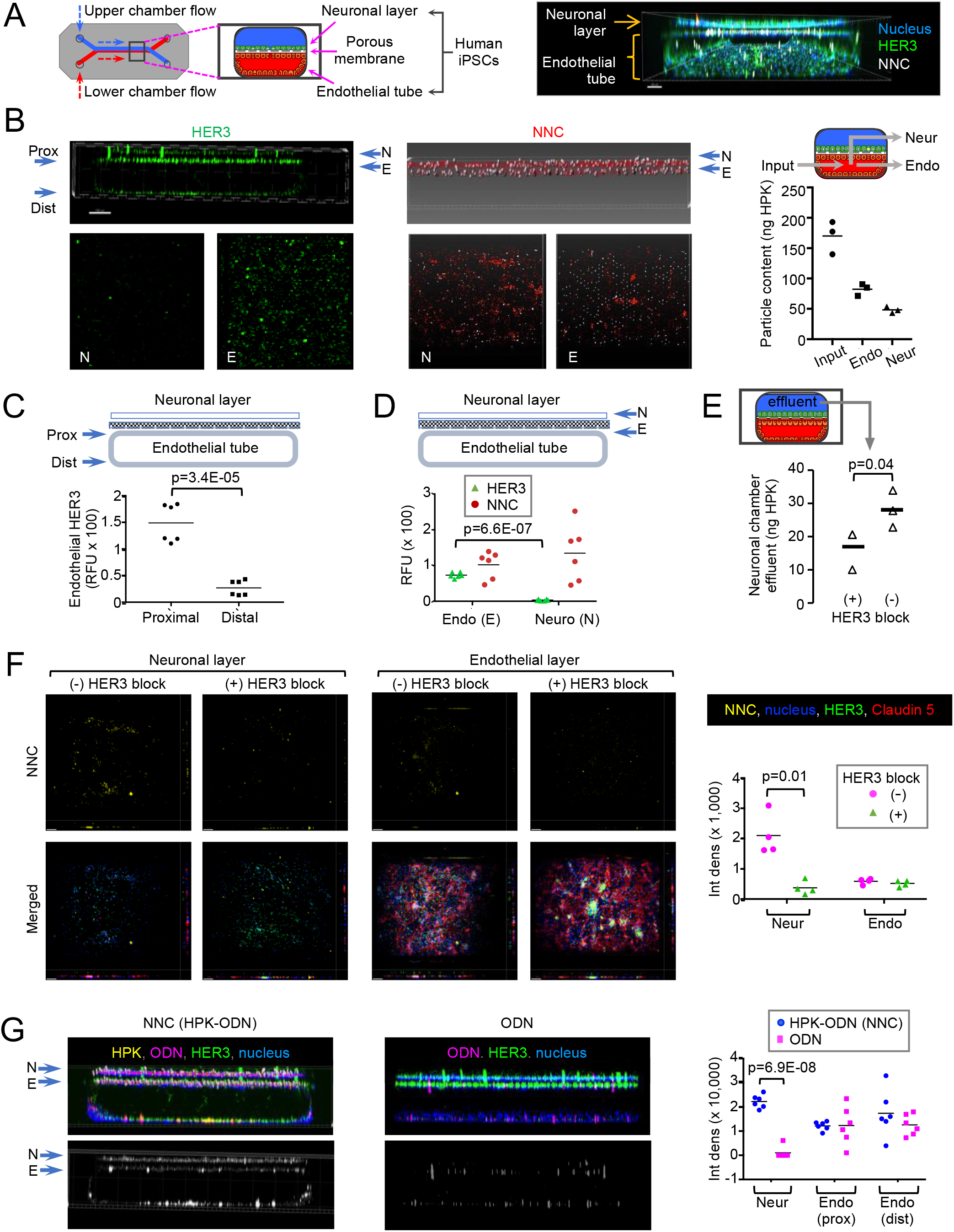
NNC transit in a reconstituted human BBB. **A,** Left, schematic of BBB chip architecture showing endothelial and neuronal chambers separated by porous membrane, and micro-ducts directing flow into and out of each chamber. Right, 3D imaging of chip at 24h after introduction of NNCs into the endothelial flow chamber. Scale bar, 150 μm. **B,** Detection of HER3 and NNCs in BBB chips at 4h after injection of NNCs delivering NIR-ODN into endothelial flow chamber. Images show cross-sectional and 2D surface views of neuronal (N) and endothelial (E) layers. Membrane pores are visible as regularly-spaced white puncta. Scale bar, 150 μm. Graph, quantification of NNCs collected from endothelial (Endo) and neuronal (Neur) chamber effluents during 4h of NNC flow into endothelial micro-chamber. Input, NNCs measured from injection micro-chamber. Data represent mean of triplicate sample effluents quantified by sandwich ELISA. **C,** Relative HER3 levels on proximal and distal endothelial layers from BBB chip represented in ***B***. Data represent averages of two regions of interest (ROIs) per zone per triplicate sample. **D,** Relative levels of HER3 and NNC on endothelial tube (E) and neuronal layer (N) from ***B***. Data represent averages of two ROIs per zone per triplicate sample. **E,** Quantification of NNC content collected from neuronal chamber effluents during 4h of NNC flow into endothelial micro-chamber -/+ blocking of HER3. NNCs were detected by sandwich ELISA using antibody capturing HPK. Data show mean and individual measurements from triplicate BBB chip samples. **F,** Immunofluorescence detection of NNCs in neuronal and endothelial layers after 4h of NNC flow into the endothelial micro-chamber -/+ blocking of HER3. Top panels, NNC fluorescence channel alone. Lower panels, merged fluorescence channels. Scale bar, 150 μm. Graph, integrated signal densities of NNCs in neuronal and endothelial compartments -/+ blocking of HER3. Data show the mean and individual measurements from 4 independent fields in each region. **G**, Comparison of NNCs delivering NIR-ODN compared to NIR-ODN alone. Upper panels, merged fluorescence channels. Lower panels, NIR channel. Graph, Integrated signal densities comparing NIR-ODN -/+ HPK in neuronal layer, proximal endothelium and distal endothelium. Data show the means from 6 independent fields in each region and mean of these individual means.

### NNCs accumulate at intracranial tumors after systemic delivery

Our findings thus far predict that brain extravasation of systemic HPK NNCs may facilitate entry into HER3-expressing intracranial tumors. To test this, we introduced NNCs carrying NIR-ODN (HPK-NIR-ODN) systemically by a single tail vein injection into mice bearing intracranial 4T1 TNBC tumors tagged with luciferase and GFP (4T1lucGFP) implanted in the right hemispheres (**Fig. 6A**). The same free NIR-ODN control used in the BBB chips earlier enabled us to confirm here whether there was any vascular leakage. To prevent this possibility, tumors were implanted 8 days before mice received systemic NNCs, enabling sufficient time for any damaged vessels to repair. At 4h after injection, systemic NNCs carrying NIR-ODN showed preferential accumulation at intracranial tumors compared to free NIR-ODN, which was either undetectable in the brains or showed no coincident localization with tumors (**Fig. 6B**). Closer histological examination of the latter confirmed that the free NIR-ODN was restricted to the vasculature in both tumor and non-tumor areas whereas HPK-NIR-ODN showed considerable extravascular spread throughout the tumor tissue (**Fig. 6, C-D**). HPK NNCs coincided with caveolin-positive sites along the brain-tumor vasculature (**Fig. 6E**), suggesting that extravasation may occur by caveolae-mediated transcytosis. HPK NNCs also coincided with the intracerebral TNBC cells and showed low to undetectable delivery to non-tumor tissue (liver, spleen, heart, lung) and non-tumor areas of the brain (**Fig. 6F**).

**Figure 6.**
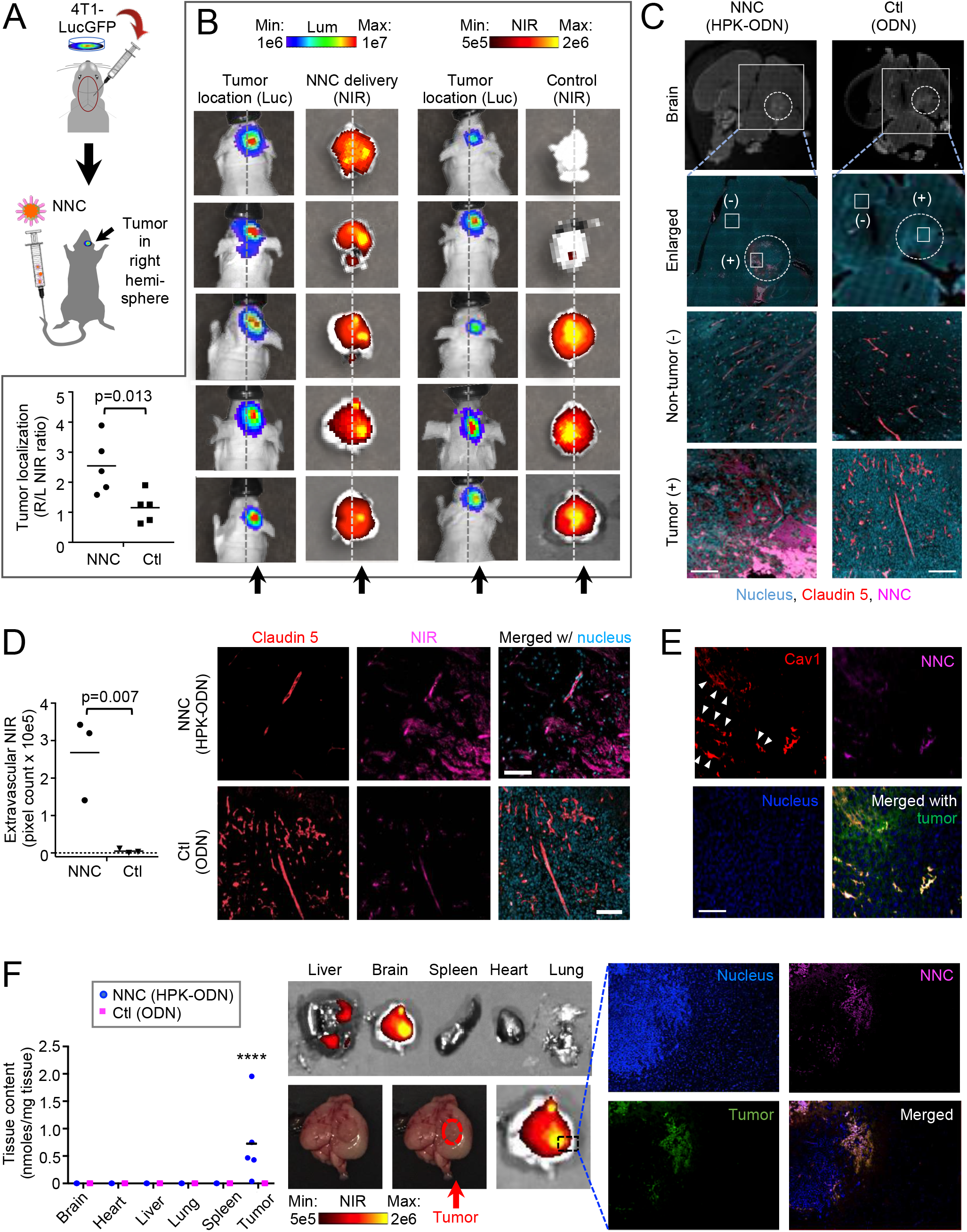
NNCs localize to intracranial tumors after systemic delivery. **A-B,** Systemic delivery of NNCs in mice bearing intracranial (IC) TNBC tumors expressing luciferase and GFP (4T1-LucGFP). Schematic in ***A*** shows the brain location of each stereotactic tumor implant. Images in ***B*** show the localization of NIR-ODN in relation to IC tumors at 4 h after tail vein injection of 1.5 nmol NIR-ODN alone or delivered by HPK NNCs. Vertical arrows point to tumor locations in right hemispheres. Left and right hemispheres are delineated by a dashed midline for reference. Graph summarizes the right/left hemisphere ratios of NIR comparing mice receiving NNCs and NIR-ODN alone. Data show mean and individual measurements of 5 mice per group. **C,** Representative brain specimens from mice in ***B*** comparing NIR fluorescence in tumor (+) and non-tumor (-) regions at 4 h after systemic delivery of NNCs or control reagents. Dashed circle delineates tumor. Scale bar, ~35 μm. **D,** Channel-separated and merged images of tumor regions shown in ***C***. Graph summarizes the quantification of extravascular (non-overlapping with claudin 5) NIR comparing specimens from mice receiving NNC or control reagents. Data show mean and individual measurements of pixel counts from 3 independent 136 μm^2^ fields all acquired at equal magnification and gain. Scale bar, ~35 μm. **E,** Specimen of IC tumor from representative mouse in ***B*** after receiving systemic NNC and showing counterstain for caveolin (Cav1). Scale bar, ~35 μm. **F,** Comparative distribution of NNCs or control reagents in tissue harvested from representative mice from ***B***. Graph summarizes the NIR measurements collected from each tissue with each point representing an individual subject (n=5). ****, p<0.0001 compared to all other tissue. Images of representative tissue are shown. Enlarged region shows brain specimen comparing localization of NNCs to tumor and adjacent non-tumor region.

### Systemic NNCs delivering chemotherapy to intracranial tumors

Taken together these findings support a mechanism in which systemically administered NNCs can cross the BBB using a HER3-mediated route associated with caveolae and accumulate in intracranial (IC) TNBC tumors using HER3 entry and subsequent pH-mediated membrane disruption (**Supplemental Fig. S5**). To examine the therapeutic efficacy of NNCs in this context, we systemically delivered HerDox in comparison to the FDA-approved liposomal doxorubicin formulation, Lipodox, at equivalent doses and regimens in mice bearing intracerebral 4T1luc/GFP tumors and monitored tumor growth by bioluminescence imaging (BLI) and magnetic resonance imaging (MRI) (**Fig. 7A and Supplemental Fig. S6**). HerDox significantly reduced and nearly flattened the average tumor growth rates as measured by BLI in contrast to mock (saline) -treated mice whereas Lipodox showed no significant difference compared to the mock-treatment (**Fig. 7B**). The broad distribution of tumor growth rates from the Lipodox-treated group also yielded no significant differences compared to the HerDox treatments based on relative tumor bioluminescence per cohort (**Fig. 7B**); however, when mice were stratified based on rates of increase in brain bioluminescence, three main phenotypes emerged that distinguished the HerDox from the Lipodox treatments. The first was characterized by a sudden exponential increase in brain bioluminescence when tumors exceeded 10^8^ relative luminescence units (RLU) that was accompanied by concomitant decline in health. 43% of the Mock cohort (6/14) and 33% of the Lipodox cohort (4/12) exhibited this characteristic whereas only 1 mouse (8% of the HerDox cohort; N=12) showed this phenotype by day 15 after tumor implant (**Fig. 7C**). The remaining mice of each cohort comprised the second phenotype characterized by luminescence retained below 1e8, which comprised 92% of the HerDox cohort compared to 57% of the Mock and 67% of the Lipodox cohorts (**Fig. 7C**). A subset of this second phenotype was characterized by negligible tumor growth and comprised 42% of both HerDox and Lipodox cohorts but only 7% of the Mock cohort (**Fig. 7C**). Tumor imaging in live mice reflected the quantified growth rates, with tumor volume differences becoming more apparent after day 11 post-implant (**Fig. 7D**).

**Figure 7.**
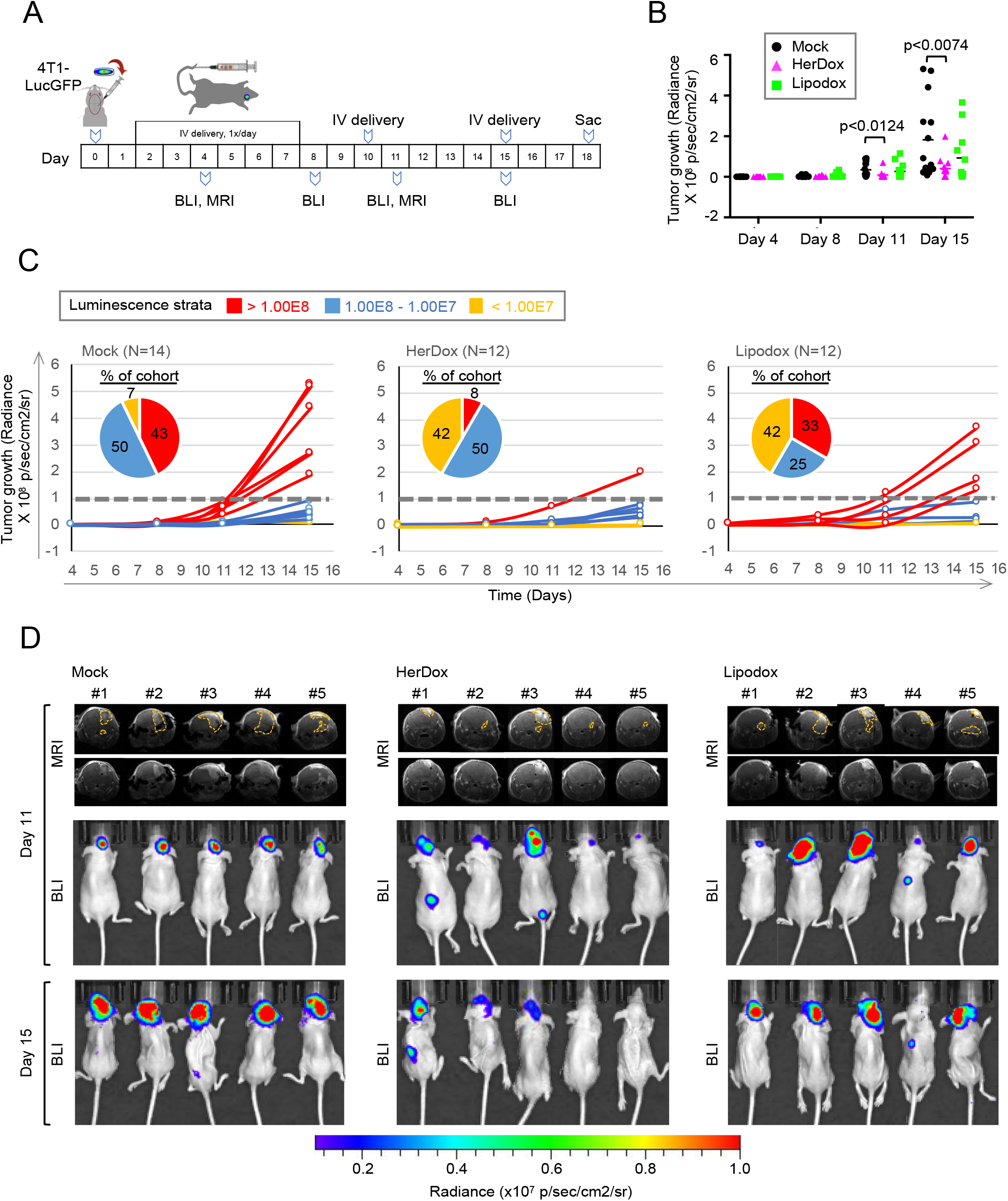
HerDox benchmarked against Lipodox in IC TNBC model. **A,** Tumor implantation, treatment and assessment regimen of mice bearing intracranial (IC) TNBC tumors. BLI, bioluminescence imaging. MRI, magnetic resonance imaging. **B-C,** Quantification of IC tumor growth. In ***B***, each data point represents the tumor radiance from individual mice of each cohort (n=12 each for HerDox and Lipodox, n=14 for Mock) with means indicated, and all measurements thresholded equally. Significances determined using 2-way ANOVA followed by Tukey post-hoc test. In ***C*,** tumor growth in each mouse is plotted for each cohort. Each growth curve is designated a color based on accelerated growth and declining health (or disease progression, shown in red), slowed tumor growth (blue), and no detectable tumor growth (or partial response, shown in yellow). **D,** MRI and BLI imaging of representative mice from each cohort. MRI images are shown with and without outlines of tumors as delineated by blinded MRI-core staff.

MRIs acquired on day 11 captured the tumor volumes mid-study (**Fig. 7D**) and showed that HerDox treatments already yielded a significant difference in size compared to the Mock treatment group (**Fig. 8A**). Although the Lipodox-treated cohort showed no significant differences in MRI-detected tumor volumes compared to the Mock and HerDox -treated cohorts at this time point (**Fig. 8A**), representative members of the Lipodox-treated group showed worsened health deterioration including extracranial metastases and reduced ambulatory ability whereas such characteristics were not detectable in the HerDox-treated mice at this time point (**Fig. 8B and Supplemental Movies 2-3**). These observations compelled us to sacrifice a representative group of mice from each cohort on day 15 (after the final bioluminescence imaging) to perform comparative tissue and blood analyses while monitoring remaining mice to assess whether similar differences between the cohorts continued (**Table 1**). The aforementioned differences in mouse health and mobility were reflected by the considerably higher proportions of remaining Lipodox (5/7) and Mock (10/10) -treated mice that required early euthanasia due to emaciation and related deteriorating body condition (Body Condition Score or BCS of less than 2)^54-55, 63^ (**Table 1**) compared to the HerDox-treated group (3/7) despite the similar decline in weights between all cohorts toward the end of the study (**Fig. 8C**). Likewise, although blood analytes signifying organ toxicity (ALT, AST, BUN, creatinine) did not significantly differ between cohorts (**Table 2**), a higher percentage of HerDox-treated mice (4/7) avoided early euthanasia compared to the Lipodox-treated mice (2/7) and in contrast to the Mock-treated cohort (0/10) (**Fig. 8D**).

**Figure 8.**
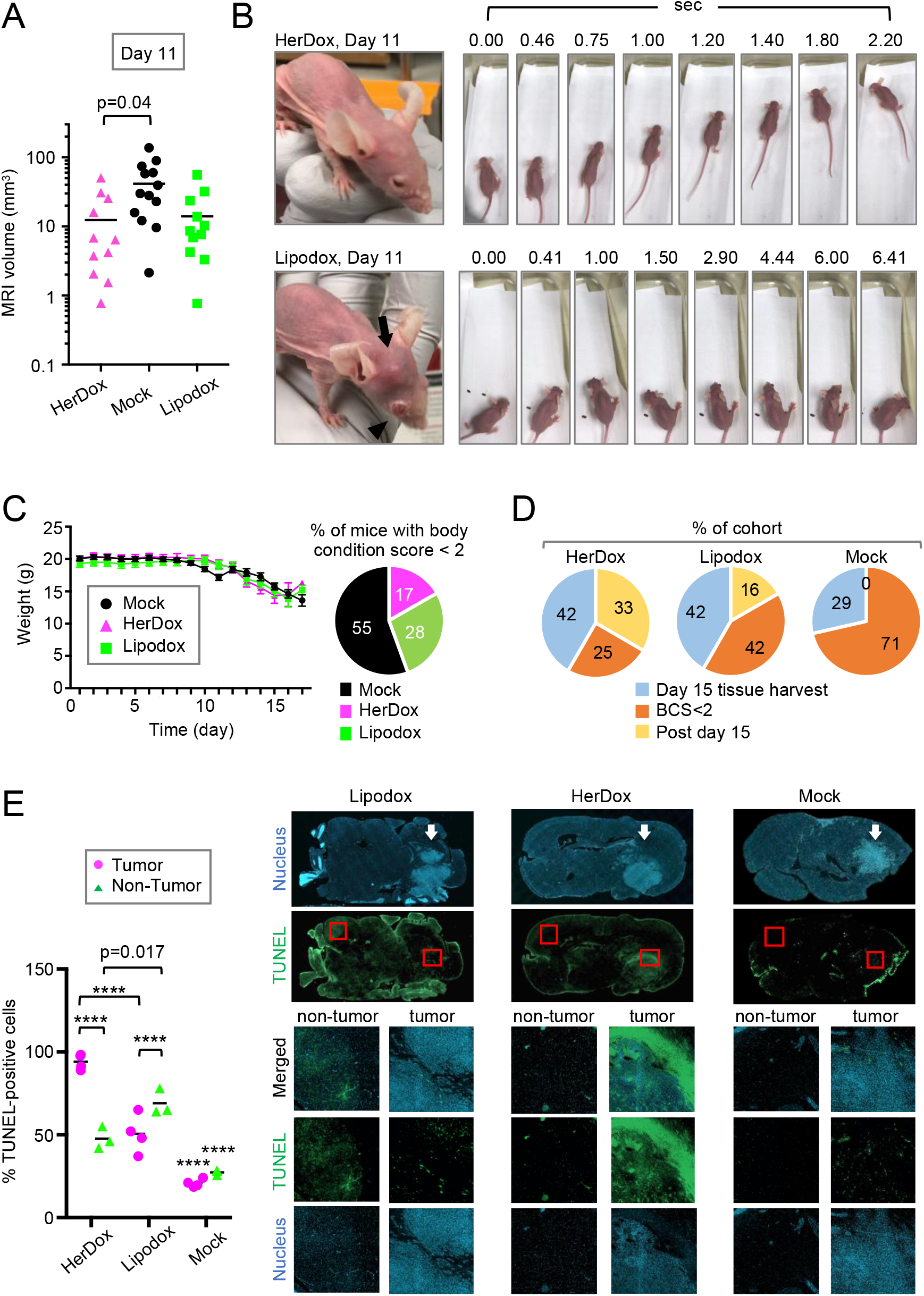
Toxicity profile in IC TNBC model. **A,** Quantification of IC tumor volumes determined by MRI at mid-treatment (Day 11 of regimen shown in ***Fig. 7A***). Data show mean and individual measurements of each cohort. **B,** External health and motility of representative mice from HerDox and Lipodox -treated cohorts on Day 11. Arrow points to extracranial tumor growth with disease affecting the eye (arrowhead). Image stills are extracted from videos in ***Supplemental Movies 2-3***. **C,** Average weights of each cohort (mean±SD) throughout experimental timeline. Pie chart summarizes the breakdown of mouse numbers requiring euthanasia due to low (<2) body condition score (BCS). **D,** Breakdown of mouse numbers reflecting the percentage of each cohort undergoing euthanasia or natural death due to the indicated circumstances. Post day 15 describes mice who survived past day 15 but eventually succumbed to disease by the endpoint. **E,** Detection of apoptosis by TUNEL stain in brain specimens from representative mice of each cohort. Channel separated images show tumor and non-tumor areas of brain specimens from each cohort. Graph summarizes the quantification of TUNEL positive cells in tumor and non-tumor areas from each cohort. Data are shown as mean and each data point indicating the percentage of TUNEL-positive cells in 3 independent fields (each averaging 227 cell events) in each indicated region and treatment. ****, p<0.0001.

**TABLE 1.** Number breakdown of treated mice bearing IC tumors.

|  | Mock | HerDox | Lipodox |
| --- | --- | --- | --- |
| N <sup>†</sup> | 14 | 12 | 12 |
| Day 15 tissue harvest <sup>‡</sup> | 4 | 5 | 5 |
| BCS < 2 <sup>§</sup> | 10 | 3 | 5 |
| Post Day 15 <sup>#</sup> | 0 | 4 | 2 |

<sup>†</sup> Total per cohort.
<sup>‡</sup> Sacrificed after final BLI for tissue/blood harvest and comparative assessments.
<sup>§</sup> Requiring euthanasia before endpoint due to no/low mobility and a Body Condition Score of less than 2 (emaciation, prominent skeletal structure, little/no flesh cover, visible and distinctly segmented vertebrae).
<sup>#</sup> Surviving beyond final BLI without requiring euthanasia.

**TABLE 2.** Blood analytes from treated mice bearing IC tumors. ^†^.

|  | Mock | HerDox | Lipodox |
| --- | --- | --- | --- |
| AST (U/L) | 1382 ± 545.1<sup>ns</sup> | 704.3 ± 310.4<sup>ns</sup> | 328.5 ± 36.9<sup>ns</sup> |
| ALT (U/L) | 209.3 ± 110.5<sup>ns</sup> | 62.3 ± 19.8<sup>ns</sup> | 21.2 ± 7.3<sup>ns</sup> |
| BUN (mg/dL) | 24.4 ± 3.9<sup>ns</sup> | 26.7 ± 6.1<sup>ns</sup> | 14.1 ± 6.3<sup>ns</sup> |
| Creatinine (mg/dL) | 0.54 ± 3.7<sup>ns</sup> | 0.8 ± 0.4<sup>ns</sup> | 1.3 ± .76<sup>ns</sup> |

<sup>†</sup> N=3 mice of each cohort.
<sup>ns</sup> No significance, determined by one-way ANOVA followed by Tukey post-hoc test. Values reflect mean +/- SEM.

Harvested brains of Lipodox-treated mice showed significantly higher damage to non-tumor areas of the brain in contrast to HerDox when brains were analyzed for apoptotic cell death (TUNEL stain; **Fig. 8E**). A comparison of tumor vs non-tumor areas showed that HerDox yielded significantly higher TUNEL-positive tumor cells compared to the non-tumor area of the same brain (**Fig. 8E**). Additionally, HerDox tumors showed significantly higher TUNEL-positivity compared to Lipodox tumors (**Fig. 8E**). In contrast, Lipodox elicited significant TUNEL-positivity in non-tumor areas throughout the brain in contrast to HerDox which showed significantly lower and less detectable TUNEL-positivity in non-tumor areas of the brain (**Fig. 8E**). Taken together these findings suggest that HerDox retains the tumoricidal activity of doxorubicin while providing more targeted toxicity to brain-localized tumors and improved preservation of healthy brain tissue compared to Lipodox.

## DISCUSSION

Here we have demonstrated that systemically administered bioparticles carrying nucleic acid cargo, or nano-nucleocapsids (NNCs) can cross the BBB and accumulate in intracranial (IC) TNBC tumors in mice using the resistance and metastasis receptor, HER3, to mediate both routes. Systemic NNCs containing intercalated doxorubicin, or HerDox, reduced growth of IC TNBC tumors in mice while having low to negligible apoptotic effect on normal brain tissue in contrast to clinically-used liposomal doxorubicin (Lipodox) which represents the currently available clinical recourse for IC tumors. We show that HER3 is prominently expressed on both healthy adult mouse and human brain endothelium while the brain parenchyma of both species exhibit low to negligible HER3. Moreover, using a human iPSC-derived BBB chip we show that NNCs recognize HER3 on the human blood vessel wall and transit across the human BBB using HER3-mediated transport.

This study demonstrates that HER3 can be used as a new and singular route of nano-carrier transfer across the BBB and into IC tumors. The association of HER3 with brain metastases and brain-localized tumors has gained increasing evidence in the oncology field ^14-28, 34-36^. Although the ErbB gene family is associated with neural growth and development,^46^ the presence of ErbB family members on the BBB has received less attention. One exception is a study by Kastin et al. (2004) who used radiolabeled neuregulin-1-ß1 to show that blockade of HER3 and HER4 but not HER2 prevents entry of systemic neuregulins into the brain parenchyma ^64^. That study suggests that HER3 normally functions on the brain endothelium to transcytose neuregulins from the circulatory system. In contrast HER2, while present on the brain endothelium^5^, does not transcytose across the blood vessel wall,^6–7^ which may explain the inability of HER2 antibodies to breach the BBB. Taken in its larger context, this suggests that the presence of a receptor on the BBB does not necessarily ensure passage of anti-receptor antibodies across the BBB and highlights the utility of ligand-mimicking molecules such as HPK that can overcome such limitations. In particular, HPK would have distinct advantages over targeting “traditionally” accepted BBB receptors such as the transferrin and glucose receptors (TfR, GLUT1). Bioinformatics analyses here suggest that HER3 is expressed at significantly higher levels on brain-metastatic breast tumors compared to TfR and GLUT1, while showing significantly lower expression than TfR and GLUT1 in normal peripheral tissue including non-CNS endothelia, thus augmenting the potential efficacy of using HER3 to target IC tumors.

The ligand-mediated approach used here for targeting across the BBB and into IC tumors offers advantages compared to antibody-based targeting. HPK used in this context is based on the premise that circulating neuregulins would require transport across the endothelium through a non-degradative pathway in order to be released into the brain parenchyma undigested ^65^. Transcytosis across the endothelium is typically mediated through caveolae trafficking,^66^ which is distinct from classical clathrin-mediated endocytosis that sorts vesicles to a lysosomal degradation pathway after transitioning through vesicle acidification.^67^ In agreement, our data show that caveolin coincides with vascular-associated NNCs in the brain which in turn emerge into the parenchyma with the cargo; thus such a pathway could support the unperturbed passage of NNCs across the BBB. HER3 undergoes a dramatic conformational change upon ligand binding ^68^, triggering ligand uptake and trafficking. This is evident in our previous studies showing that HPK particles rapidly sequester HER3 to clustered sites at the surface of tumor cells in contrast to resting tumor cells and these HER3 clusters coincide with particle uptake and trafficking occurring within minutes after cell binding ^42, 44^. This contrasts with monoclonal antibodies and antibody-drug conjugates that produce negligible to undetectable uptake or comparatively slow (up to 24h or more) and passive uptake after binding to target cells.^69–71^ Meanwhile, we have observed here and in earlier studies^42, 44, 49^ that HPK particles lacking any drug show no growth-stimulating effect and may in some circumstances contribute to reducing tumor growth. The latter may be attributed in part to the sequestration of HER3 by the multivalency of the particle^44^ and by the ability of the HPK ligand to reduce HER3 activation by wild-type heregulins.^42^

While most ADCs lack an endosomolytic domain, other types of nano or bio-carriers may include peptides or devices intended to provide an endosomal escape function after cell uptake. ^72–73^ Too often, sufficiently rigorous mechanistic studies are lacking in empirically demonstrating the utility of these moieties ^72^. These latter concerns are among those raised in light of the numerous antibody-drug conjugates that have been produced for the clinic but have generally yielded weak clinical outcomes.^74^ Accordingly, we continue to probe and add to our understanding the elusive mechanism underlying the endosomolytic function of the penton base capsomere and its derivative proteins such as HPK. We have previously found that the solvent accessible pore comprising the capsomere barrel is lined with histidines and charged residues whose protonation should cause repellence of HPK monomers and exposure of hydrophobic domains mediating pentamerization ^75^. Here we provide additional evidence that HPK monomers extend away from the capsomere axis under increasing acidification conditions using molecular dynamics simulation, supporting our earlier functional studies showing that capsomeres reduce to smaller constituents under decreasing pH ^75^. We also show here that HPK avoids distribution to Rab7+ endo-lysosomes and enters the cytoskeletal compartment in contrast to the ligand lacking the penton base domain, which remains associated with the membrane compartment after uptake. Cytoskeletal attachment facilitates the post-endosomal trafficking of adenovirus ^76–77^ and its capsid proteins ^47, 78^ inside the cell. Taken together, we propose that the exposure of hydrophobic pentamerization domains upon low pH may enable the disruption of vesicle lipids, thus weakening the endosome wall. In agreement, we observe here that inhibiting vesicle acidification retains HPK capsomeres in vesicle-like puncta and reduces diffusion into the extravesicular space. This naturally evolved structure afforded by the penton base domain provides a potentially modulatable function enabling low pH-mediated penetration into tumor cells that avoids premature release in the endothelium as shown in this study.

Use of the human iPSC-derived BBB chip validated our findings in mice, showing that HER3 is prominently expressed on the human adult BBB in contrast to non-BBB endothelia and extravascular brain parenchyma; and that NNCs use HER3 to extravasate the brain vasculature. The cross-reactivity of NNCs to both mouse and human HER3 enable pre-clinical translational value to be drawn from both our human tissue and *in vivo* mouse studies. Specifically, lack of access to intravital imaging limited our mouse studies to using *in vivo* and *in situ* detection and immunofluorescence analyses for examining brain extravasation in mice; however, these findings are supplemented by studies in human BBB chips which not only allow us to validate our BBB extravasation findings in a dynamic model but also to test this in a human system. The benefit of species cross-reactivity further supported our therapeutic efficacy studies in which benchmarking the HerDox NNCs against Lipodox was tested in a syngeneic 4T1 model mimicking late stage TNBC. While these studies used immunodeficient instead of immunocompetent mice, the immune deficient setting reduced chances of spontaneous tumor rejection and enabled sufficient tumor growth (with 100% IC tumor take) to provide the background against which HerDox performance can be assessed. Although immunogenicity assessments are lacking in this study we have previously performed immunostimulatory evaluations in similar studies using NNCs to deliver siRNA (designated HerSi) in BALB/c mice ^44^. Those studies showed that repeat inoculations of HerSi at 10 times the dosage used for therapeutic efficacy yielded no significant increase in serum Ig levels in contrast to adenovirus used as an immunostimulatory control ^44^. Moreover, we have shown in those studies that antibodies recognizing the penton base are partially masked from immunorecognition in the context of the HPK capsomere and fully masked in the assembled bioparticle ^44^.

In conclusion, these studies demonstrate that HER3 is expressed on the blood-brain barrier and mediates extravasation and tumor-targeting of systemic ligand-mimicking bioparticles for reducing growth of intracranial triple-negative breast cancer. Given the growing range of tumor types showing association of HER3 with resistance and metastasis, these studies can pave the way for using HER3 bioparticles to target such tumors when localized in the brain.

## ACKNOWLEDGMENTS

The authors thank the Cedars-Sinai Research Imaging core for animal imaging; the Electron Imaging Center for NanoMachines (EICN) at UCLA for EM services; the ICP-MS core at UCLA for quantification of metal content in tissues; Zeev Gross, PhD and Harry Gray, PhD for use of gallium-metallated corroles in the biodistribution study using ICP-MS; Omar Haffar and Kent Iverson for critical review and intellectual feedback on this work; and BioScience Writers for editorial assistance. LMK thanks C Rey and AACM Kauwe for ongoing support.

## FUNDING

This research was supported by grants from the National Institutes of Health (NIH): NCI R01 CA129822, R01 CA140995, and NCATS UL1 TR001881; and the Department of Defense (DoD): BCRP W81XWH-15-1-0604, W81XWH1910592. F.A.-V. was supported in part by a training grant from National Institutes of Health [T32 HL134637].

## SUPPLEMENTAL MATERIALS

### SUPPLEMENTAL METHODS

ITC of HerDox

Therapeutic efficacy in mice with peripheral tumors

### SUPPLEMENTAL FIGURES

Supplemental Figure S1. Immunoblots of post-endosomal fractions.

Supplemental Figure S2. HPK capsomere structure and HerDox particle assembly.

Supplemental Figure S3. Systemic HerDox compared to Lipodox on BALB/c-4T1 peripheral tumors.

Supplemental Figure S4. Tissue distribution and time course.

Supplemental Figure S5. Summary of HER3-mediated BBB passage and tumor entry.

Supplemental Figure S6. Bioluminescence imaging of mice at indicated days of IC tumor growth.

### SUPPLEMENTAL MOVIES

Supplemental Movie 1. HPK capsomere undergoing protonation. https://www.youtube.com/watch?v=QiX_6bY_hig

Supplemental Movie 2. Representative mouse from HerDox cohort. https://www.youtube.com/watch?v=ORx8oWy9H_w

Supplemental Movie 3. Representative mouse from Lipodox cohort. https://www.youtube.com/watch?v=QZNHg3rXzt0

## Notes

### Competing Interest Statement

L.K. Medina-Kauwe and Cedars-Sinai Medical Center hold significant financial interest in Eos Biosciences, Inc., of which L.K. Medina-Kauwe is co-founder and scientific advisor.

