## Supplemental materials for "Systemic ligand-mimicking bioparticles cross the blood-brain barrier and reduce growth of intracranial triple-negative breast cancer using the human epidermal growth factor receptor 3 (HER3) to mediate both routes"

**Contents**

***SUPPLEMENTAL METHODS***

ITC of HerDox

Therapeutic efficacy in mice with peripheral tumors

***SUPPLEMENTAL FIGURES***

Supplemental Figure S1. Immunoblots of post-endosomal fractions.

Supplemental Figure S2. HPK capsomere structure and HerDox particle assembly.

Supplemental Figure S3. Systemic HerDox compared to Lipodox on BALB/c-4T1 peripheral tumors.

Supplemental Figure S4. Tissue distribution and time course.

Supplemental Figure S5. Summary of HER3-mediated BBB passage and tumor entry.

Supplemental Figure S6. Bioluminescence imaging of mice at indicated days of IC tumor growth.

***SUPPLEMENTAL MOVIES***

Supplemental Movie 1. HPK capsomere undergoing protonation. <https://www.youtube.com/watch?v=QiX_6bY_hig>

Supplemental Movie 2. Representative mouse from HerDox cohort. <https://www.youtube.com/watch?v=ORx8oWy9H_w>

Supplemental Movie 3. Representative mouse from Lipodox cohort. <https://www.youtube.com/watch?v=QZNHg3rXzt0>

***SUPPLEMENTAL METHODS***

**ITC of HerDox**. The two-step assembly of HerDox requires two separate ITC measurements: 1) binding of Dox to the oligonucleotide duplex, dsLLAA; and 2) binding of HPK to Dox-intercalated dsLLAA. The complementary oligonucleotide duplexes forming dsLLAA were prepared by mixing together equal molar concentrations of the 30-base oligonucleotide, LLAA-5 (5’CGCCTGAGCAACGCGGCGGGCATCCGCAAG-3’), and its corresponding reverse complement LLAA-3 in annealing buffer pH 7.4. The mixture was boiled in a beaker filled with water for 5 min, then the beaker was transferred to the benchtop and allowed to cool to room temperature. All following dilutions of dsLLAA or Dox were done in Sodium Phosphate buffer (pH 7.4) to match the storage buffer of HPK.

**ITC #1) To determine Dox: dsLLAA binding molar ratio.** The ITC experiment was carried out using MicroCal PEAQ-ITC at 25 °C. 300 μL of 4.0 μM dsLLAA in the cell was titrated with 500 μM Dox. Titrations took place by injecting 2 μL Dox in a 2.5 min injection for the titration peak to return to the baseline. The K_d_ was calculated using the MicroCal PEAQ-ITC analysis software using the one-site model. Control experiments were carried out by titration of 500 μM Dox into buffer (Sod. Phosphate pH 7.4), buffer into 4.0 μM dsLLAA, and buffer into buffer. The three controls were used a composite for the ITC experiment to subtract the heat of dilution and background noise from the baseline.

**ITC #2) To determine the dissociation constant and HPK: Dox-dsLLAA binding molar ratio.** A fresh batch of Dox-dsLLAA was mixed in a molar ratio of 7: 1 Dox: dsLLAA (based on findings from ITC #1) To a final dsLLAA concentration of 12 μM. The mixture was incubated at room temperature for t least 30 minutes. The ITC experiment was carried out using MicroCal PEAQ-ITC at 25 °C. 300 μL of 3.5 μM HPK in the cell was titrated with 12 μM dsLLAA in the form of Dox-dsLLAA. Titrations took place by injecting 2 μL Dox in a 2.5 min injection for the titration peak to return to the baseline. The K_d_ was calculated using the MicroCal PEAQ-ITC analysis software using the one-site model. Control experiments were carried out by titration of 12 μM Dox-dsLLAA into buffer (sod. Phosphate pH 7.4), buffer into 3.5 μM HPK, and buffer into buffer. The three controls were used a composite for the ITC experiment to subtract the heat of dilution and background noise from the baseline.

**Therapeutic efficacy in 4T1 peripheral tumor model**. Female BALB/c mice (~6 weeks; N=5 per treatment group) received bilateral flank injections of 4T1lucGFP tumor cells (1e5 cells/implant). Approximately five days later when tumors were palpable, the mice were randomized and began receiving daily tail vein injections of HerDox or Lipodox at either 0.2 or 0.02 mg/kg doxorubicin per dose, empty particles (lacking the doxorubicin) equating the 0.2 mg/kg dose, or saline at equivalent volume for five days followed by twice weekly injections for four weeks while tumor growth was monitored by bioluminescence imaging (BLI) and manual measurement of primary tumor volume by calipers. To ensure rigor, measurements were collected in blinded fashion with cohort identities unknown to the researcher.

***SUPPLEMENTAL FIGURES***

**Supplemental Figure S1. Immunoblots of endosomal and post-endosomal fractions.** Full immunoblots of endosomal (membrane) and post-endosomal (cytosolic, cytoskeletal) fractions isolated from HER3+ MDA-MB-435 human tumor cells harvested and processed at the indicated time points during uptake of HPK or the PB-deleted construct, HΔPK. Membrane fractions are delineated by TIM23, cytosolic fractions are delineated by GAPDH, and cytoskeletal fractions are delineated by β-actin. Low MW (25 kDa) bands seen in the membrane fractions are likely produced by residual heregulin bound to the cell membrane. The MW’s of the standard ladder bands are shown in kDa.

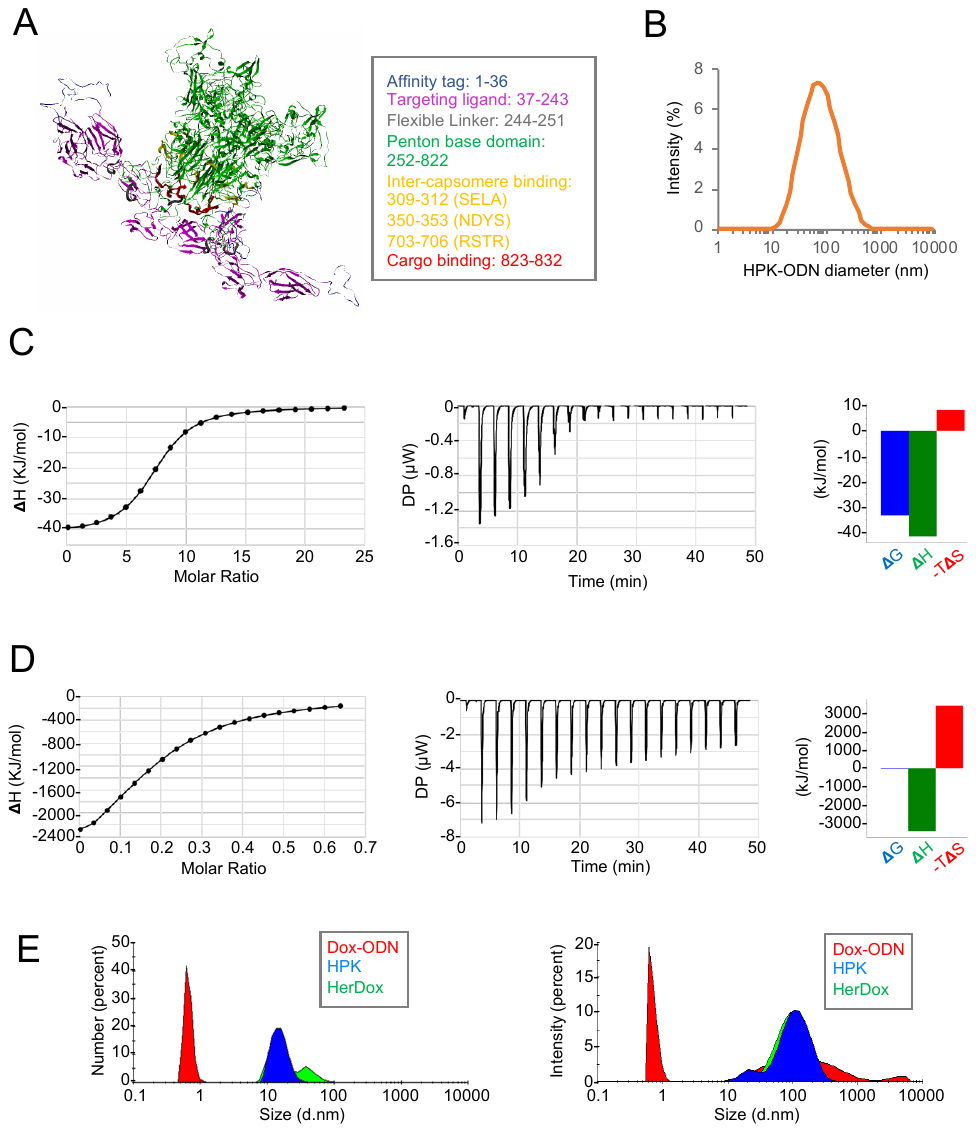

**Supplemental Figure S2. HPK capsomere structure and HerDox particle assembly.** **A**, Computational model of HPK pentamer shown as a ribbon structure with functional domains delineated by the indicated assigned colors. **B**, Intensity based particle size determination parameter from dynamic light scattering measurements of HPK+ODN. **C,** Dox binding to dsODN. Panels shown from left to right represent: the binding isotherm from the integrated thermogram fit using the one-site model in the PEAQ-ITC software (N = 7.18 ± 0.19, and K_d_ = 1.53 ± 0.39 μM), the raw thermogram used to generate the binding isotherm, and signature plot showing the thermodynamics parameters (ΔH = -41.5 ± 1.91 kJ/mol, ΔG = -33.2 kJ/mol, and -TΔS = -8.30 kJ/mol). **D**, Dox-dsODN binding to HPK. Panels shown from left to right represent: the binding isotherm from the integrated thermogram fit using the one-site model in the PEAQ-ITC software [N = 0.18 ± 0.02 (or molar ratio for HPK: Dox-dsLLAA = 5.5 ± 1.0), and K_d_ = 0.31 ± 0.15 μM], the raw thermogram used to generate the binding isotherm, and signature plot showing the thermodynamics parameters (ΔH = -3410 ± 838 kJ/mol, ΔG = -37.2 kJ/mol, and -TΔS = 3370 kJ/mol). **E**, Dynamic light scattering of HerDox and pre-assembly constituents showing particle size determination by number (left) and intensity (right).

**Supplemental Figure S3. Systemic HerDox compared to Lipodox on BALB/c-4T1 peripheral tumors. A,** Growth of 4T1 bilateral flank tumors (determined by volume measurement) in BALB/c mice during systemic treatment with indicated doses of Lipodox, HerDox, Empty (drug lacking) HPK particles (at doses equating 0.2 mg/kg HerDox) and saline. Data represent mean±SD of 5 mice per treatment group. Due to the multiple significant differences detected between treatments at specified time points (&, Day 10; #, Day 14), comparisons are shown next to the figure legend. *, p<0.05; **, p<0.01; ***, p<0.001; ****, p<0.0001. Day 0 corresponds to first day of treatment (~5 days after tumor implant). Methodological details provided in the ***Supplemental Methods***. **B,** Bioluminescence imaging of mice representing each treatment cohort. **C,** Mouse weights during indicated treatments. Data represent mean±SD of 5 mice per treatment group.

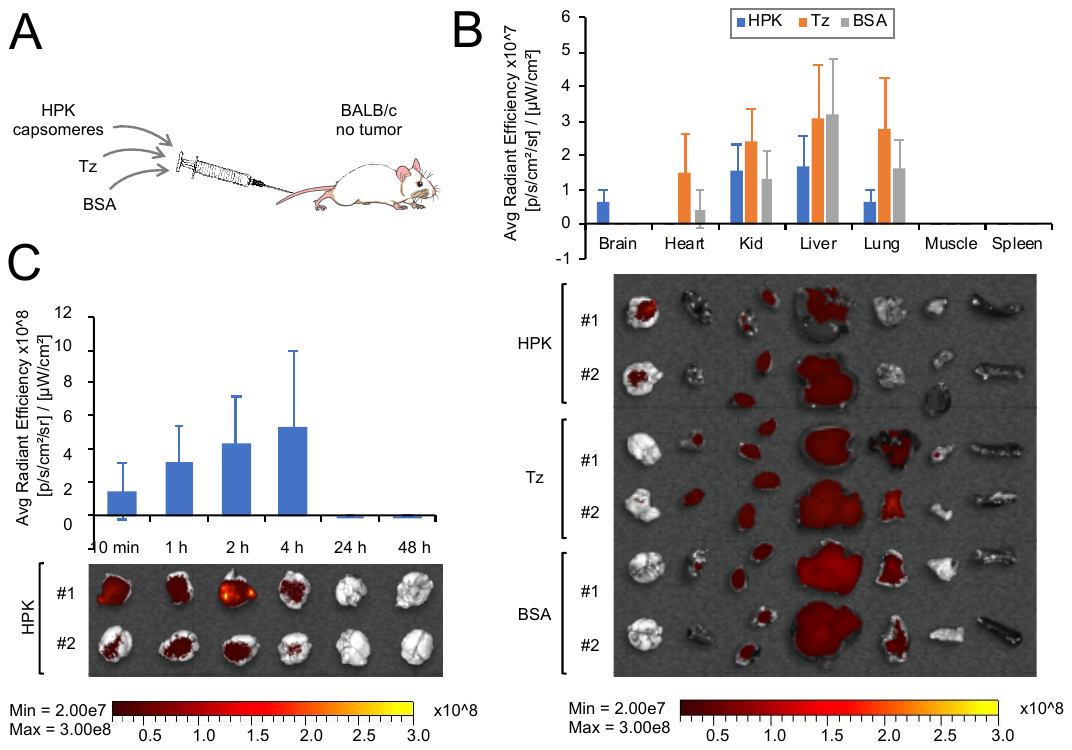

**Supplemental Figure S4. Tissue distribution and time course. A,** Schematic summarizing the conditions for systemic delivery of indicated reagents in tumor-free BALB/c mice (n=5 mice per treatment reagent). Each reagent was directly labeled with a near infrared (NIR) tag, purified and quantified before tail vein injection (12 nmol/injection). Each mouse received a single injection before sacrifice and tissue harvest at the indicated time points detailed in ***B*** and ***C*** after injection. **B,** Image acquisition and measurement of average radiant efficiencies collected from tissue harvested >4h after systemic delivery of indicated reagents as described in ***A***. Tissues shown below graph were acquired from representative mice of each cohort. **C,** Image acquisition and measurement of average radiant efficiencies collected from brains harvested at indicated time points after systemic delivery of labeled HPK capsomeres as described in ***A***. Brains shown below graph were acquired from representative mice of each cohort.

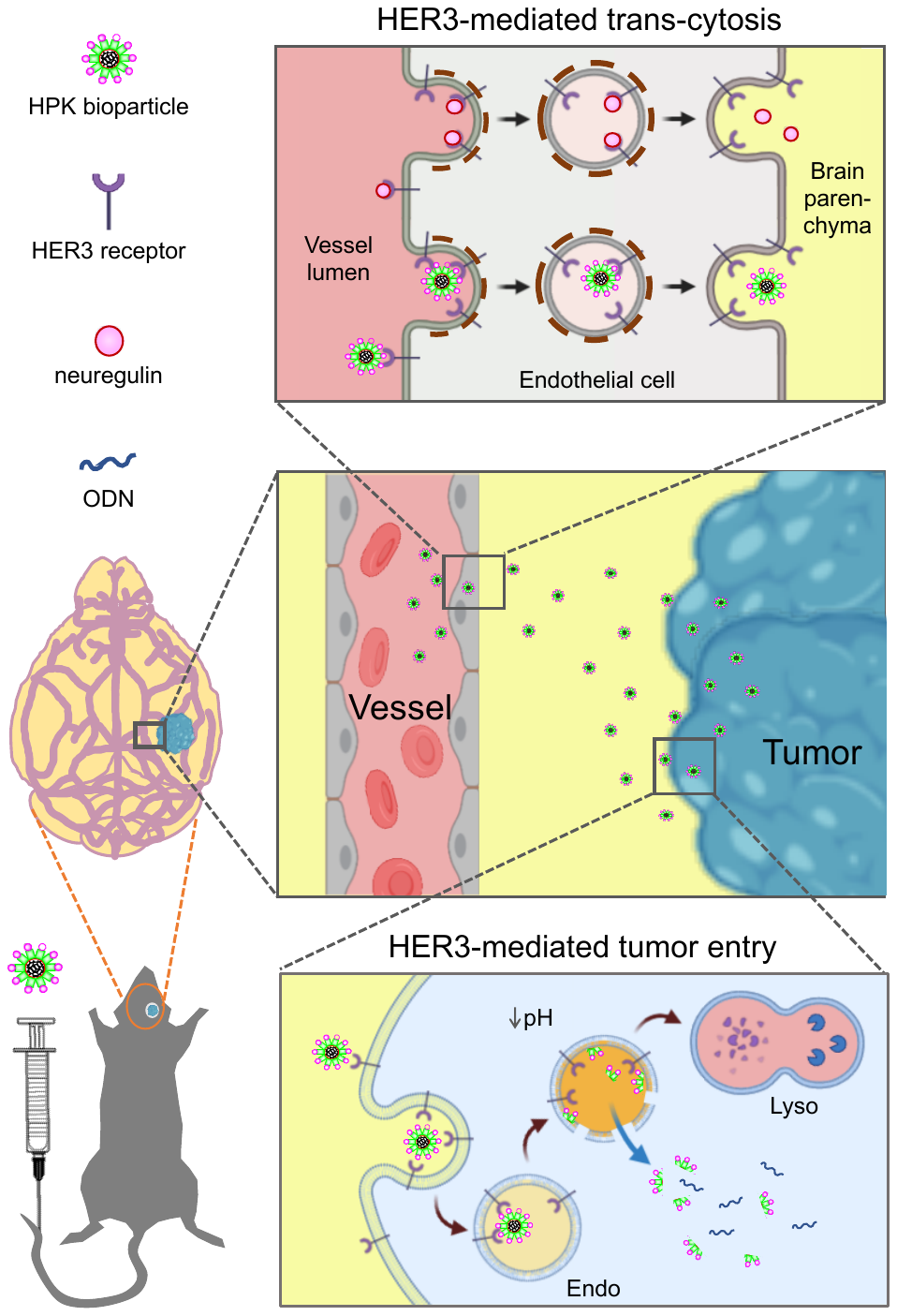

**Supplemental Figure S5. Summary of HER3-mediated BBB passage and tumor entry.** The HPK bioparticle exploits the native transcytosis pathway of neuregulin by binding HER3 receptors in the vessel lumen and undergoing non-acidifying transcytosis, followed by HER3-mediated endocytosis in tumor cells. Endosome acidification triggers opening of the HPK capsomere enabling penton base -mediated membrane destabilization and release of vesicle contents.

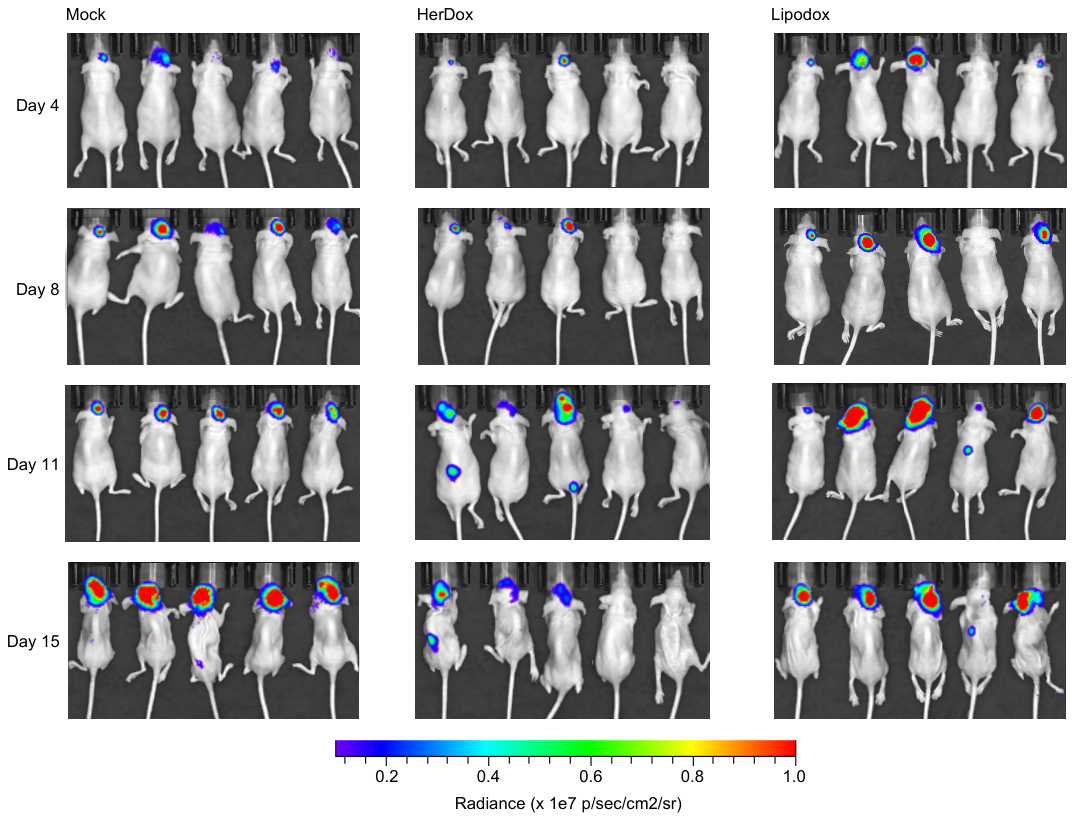

**Supplemental Fig. S6. Bioluminescence imaging of mice at indicated days of IC tumor growth.**

Bioluminescence imaging of representative mice on indicated days after implant of intracranial 4T1 tumors during indicated treatments (Mock, HerDox, or Lipodox) as described in ***Figure 7***. Each mouse was positioned similarly for each timepoint with image thresholding kept constant between cohorts, treatments, and timepoints to acquire comparative bioluminescence measurements after injections of D-luciferin (described further in Luciferase Imaging methods). Scale bar reflects the radiance from 1e6 p/sec/cm^2^/sr to 1e7 p/sec/cm^2^/sr.
